# Microtubule binding-induced allostery promotes LIS1 dissociation from dynein prior to cargo transport

**DOI:** 10.1101/2022.11.08.515461

**Authors:** William D. Ton, Yue Wang, Pengxin Chai, Cissloyny Beauchamp-Perez, Nicholas T. Flint, Lindsay G. Lammers, Hao Xiong, Kai Zhang, Steven M. Markus

## Abstract

The lissencephaly-related protein LIS1 is a critical regulator of cytoplasmic dynein that governs motor function and intracellular localization (*e*.*g*., to microtubule plus-ends). Although LIS1 binding is required for dynein activity, its unbinding prior to initiation of cargo transport is equally important, since preventing dissociation leads to dynein dysfunction. To understand whether and how dynein-LIS1 binding is modulated, we engineered dynein mutants locked in a microtubule-bound (MT-B) or -unbound (MT-U) state. Whereas the MT-B mutant exhibits low LIS1 affinity, the MT-U mutant binds LIS1 with high affinity, and as a consequence remains almost irreversibly bound to microtubule plus-ends. We find that a monomeric motor domain is sufficient to exhibit these opposing LIS1 affinities, and that this is an evolutionarily conserved phenomenon. Three cryo-EM structures of dynein with and without LIS1 reveal microtubule-binding induced conformational changes responsible for this regulation. Our work reveals key biochemical and structural insight into LIS1-mediated dynein activation.

## INTRODUCTION

Cytoplasmic dynein-1 is a highly conserved molecular motor that transports a variety of cargos toward the minus ends of microtubules in eukaryotic cells throughout the evolutionary spectrum. The dynein complex is comprised of several accessory chains (*i*.*e*., light, light-intermediate, and intermediate chains), and two copies of the approximately 500 kDa heavy chain, the latter of which possesses all of the elements required for motility^1^. Processive dynein motility requires that it first associate with its activating complex dynactin, and a cargo adaptor that link dynein to dynactin and a variety of cargoes^2,3^. Recent studies have revealed additional levels of regulation that include autoinhibitory mechanisms intrinsic to the dynein and dynactin complexes, and also for some of the cargo adaptors^2,4-8^. For example, dynein exists in an autoinhibited conformational state referred to as the ‘phi’ particle (due to its similarity to the Greek letter) that restricts its ability to bind to dynactin and the cargo adaptor^7,8^.

In addition to dynactin and cargo adaptors, several other regulators of dynein have been shown to impact its activity. Among the most important of these are the lissencephaly-related protein LIS1, mutations in which lead to dynein dysfunction and severe neurodevelopmental disorders^9,10^. Studies from several model systems including budding yeast, filamentous fungi, and humans have supported a model whereby LIS1 binds to dynein when it is in its open state, and prevents it from switching back to the phi particle^8,9,11-13^. LIS1 has also been shown to promote dynein’s association with microtubule plus ends^14-19^, and to aid in the recruitment of a second dynein complex to dynactin, thereby stimulating formation of faster motor complexes^12,13,20,21^.

Although LIS1 binding to dynein is required for it to promote these activities, several lines of evidence suggest that LIS1 dissociates from dynein prior to initiation of cargo transport. For example, LIS1 homologs in filamentous fungi were only found to transiently associate with retrograde-moving dynein-driven endosomes^22,23^. Similarly, although the LIS1 homolog Pac1 associates with dynein at the plus ends of microtubules in budding yeast, it does not colocalize with dynein at its site of activity in this organism: the cell cortex^14,24^. Studies using purified proteins revealed that LIS1 and Pac1 only associate with a small fraction of motile dynein complexes in single molecule assays^8,12,13^. Perhaps most strikingly, whereas LIS1 associates with and promotes plus end binding of dynein in reconstituted assays, only a small fraction of dynein-dynactin-adaptor (DDA) complexes had detectable LIS1 bound^25^. This is consistent with data from budding yeast in which overexpression of the dynein-dynactin-binding domain of the cargo adaptor Num1 appears to promote assembly and motility of DDA complexes that do not colocalize with Pac1^26^.

Whether dissociation of dynein-LIS1 complexes is required for proper cargo transport in metazoa is unclear. However, one piece of evidence from budding yeast suggests that their dissociation is indeed crucial. Specifically, inclusion of bimolecular fluorescence complementation (BiFC) tags on Pac1 and Dyn1 (the latter of which encodes the dynein heavy chain) leads to a situation in which these two proteins remain associated subsequent to delivery of dynein-dynactin to Num1 receptors at the cell cortex (due to the irreversible association of the two split-YFP halves)^27,28^. Whereas those cells expressing only one of the two BiFC-tagged proteins possess normal dynein function, those expressing both exhibit defects in dynein-mediated spindle positioning as severe as those lacking Dyn1, suggesting that dynein-Pac1 dissociation is critical for proper in-cell dynein activity.

Our understanding of LIS1 and Pac1 function is complicated by conflicting findings regarding these molecules’ abilities to modulate dynein’s microtubule-binding activity, thereby affecting its velocity, its force generation properties, and potentially its ability to remain associated with microtubule plus ends^29-31^. Arguments against this model include work from our lab revealing that a Pac1-bound dynein does not employ its microtubule-binding domain (MTBD) to associate with plus ends in cells^26^. We also discovered that the extent of Pac1’s ability to reduce dynein velocity *in vitro* directly scales with Pac1’s microtubule-binding^8,9^. Since Pac1 does not bind microtubules in cells, these findings force us to reevaluate whether Pac1 and LIS1 actually impact dynein mechanochemistry and/or force production. Determining whether LIS1 remains associated with motile dynein-dynactin complexes in cells, and understanding the consequences of preventing their dissociation will clarify these controversies, and ultimately reveal LIS1’s true activities.

Here we set out to address the question of whether and how dynein-LIS1 affinity may be modulated, and to specifically address whether microtubule binding by dynein may be responsible. By using a protein engineering approach, we generate dynein mutants that are constitutively locked in either a microtubule-unbound or -bound conformational state. Our data reveal that these mutants indeed reflect the native bona fide conformations of dynein in these two states, and that they have opposing affinities for LIS1. Specifically, the microtubule-unbound state of dynein exhibits significantly higher affinity for LIS1 than the microtubule-bound state. We find that the motor domain of dynein is sufficient for this behavior, and that it is conserved from yeast to humans. Cells expressing the microtubule-unbound dynein mutant exhibit robust dynein-Pac1 binding, but little unbinding, and exhibit behavior consistent with an inability of dynein to dissociate from the plus end-binding machinery, and thus the plus ends themselves. Our observations indicate that dynein must switch to a microtubule-bound conformation in order to dissociate from LIS1, which then permits the adoption of a motility-competent state of the cortical DDA complex. High-resolution cryoelectron microscopy (cryo-EM) structures reveal the structural basis for microtubule-binding-induced dissociation of dynein-LIS1. Our data are consistent with a model in which LIS1 must dissociate from dynein prior to initiation of cargo transport, and that microtubule-binding is responsible for triggering this conformational change.

## RESULTS

### Generation of constitutive microtubule-unbound and -bound dynein mutants

A previous study from our lab revealed that binding of the coiled-coil domain of the yeast cargo adaptor protein Num1 (Num1_CC_) to dynein-dynactin triggers dissociation of Pac1 from dynein, thus promoting minus end-directed motility of the motor complex^26^. However, deletion of dynein’s MTBD prevents this dissociation, suggesting that dynein must bind microtubules for this to occur. We thus sought to determine whether microtubule-binding leads to structural rearrangements sufficient to trigger dissociation of dynein from Pac1. Microtubule binding by dynein leads to a conformational change in the MTBD that is communicated to the AAA+ ring via a translation of the CC1 helix of the dynein stalk with respect to CC2, causing a change in the heptad registry of this coiled-coil (Fig. S1A)^32-35^. We hypothesized that this helix shift is the trigger that initiates a cascade of events that ultimately leads to dissociation of Pac1 from dynein.

To test this hypothesis, we employed a protein engineering strategy in which the dynein MTBD and short regions of CC1 and CC2 are replaced with a stable coiled-coil derived from an exogenous protein, seryl tRNA synthetase (SRS_CC_)^36^. By including or excluding 4 amino acids in CC1, sufficient to encode a single turn in this helix (Fig. S1B), we aimed to lock CC1 in either an up or down state, thus reflecting the microtubuleound or -unbound state, respectively. Consistent with previous work using *Dictyostelium discoideum* dynein^36^, the microtubule-unbound (MT-U) and - bound (MT-B) mutants exhibit ATPase rates that closely match that of wild-type dynein in the absence and presence of microtubules, respectively (Fig. S1C).

### Microtubule-unbound and -bound dynein mutants exhibit opposing localization behaviors in cells

The extent to which dynein and Pac1 interact governs the degree to which these proteins localize to various sites in cells (*e*.*g*., microtubule plus ends). For example, cells with no Pac1 exhibit an almost complete lack of dynein foci^14,19^, while those overexpressing Pac1 or expressing a dynein mutant with higher-than-wild-type affinity for Pac1 both exhibit a greater number of dynein (and Pac1) foci^8,24,27^. Thus, the number and brightness of dynein foci directly correlate with dynein-Pac1 affinity. Imaging cells expressing Pac1-3mCherry and either dynein^MT-U^-3YFP or dynein^MT-B^-3YFP revealed that these two mutants exhibit opposing degrees of dynein localization. Specifically, dynein^MT-U^-3YFP-expressing cells exhibit more plus end and cortical foci than wild-type cells, while only a small fraction of dynein^MT-B^-3YFP-expressing cells exhibit fluorescent foci that were significantly less bright than those in dynein^MT-U^-3YFP-expressing cells (Fig. 1A-C, non-hatched bars).

**Figure 1.**
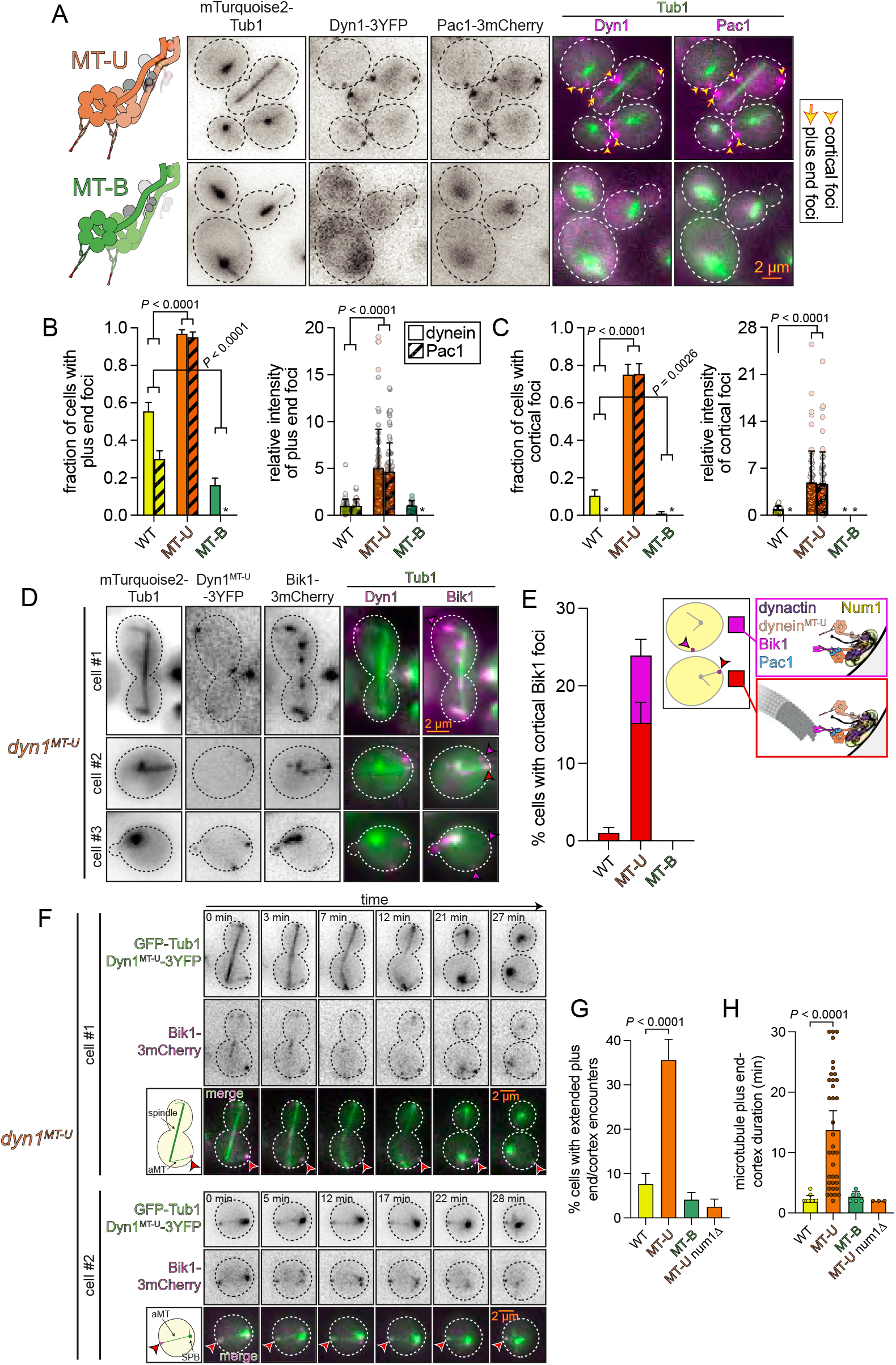
The microtubule-unbound dynein mutant is tightly bound to Pac1, Bik1, and microtubule plus ends in cells. (A) Representative fluorescence images of cells expressing Pac1-3mCherry, mTurquoise2-Tub1, and either Dyn1^MT-U^-3YFP or Dyn1^MT-B^-3YFP (arrow, plus end focus; arrowheads, cortical foci). (B and C) Plots depicting frequency and intensity of plus end (B) and cortical (C) dynein and Pac1 foci in cells expressing indicated DYN1 allele as the only source of dynein heavy chain (WT, wild-type; MT-U, microtubule-unbound mutant; MT-B, microtubule-bound mutant). Foci were scored from timelapse movies (*, no foci observed). (C) Plot depicting frequency of cells with Pac1-3mCherry foci. (D) Representative fluorescence images of cells expressing Bik1-3mCherry, mTurquoise2-Tub1, and Dyn1^MT-U^-3YFP (magenta arrowhead, cortical Bik1 foci without associated plus end; red arrowhead, cortical Bik1 focus with associated plus end). (E) Plot depicting frequency of cells with indicated dynein allele possessing cortical Bik1 foci (either with or without associated microtubule plus end, as indicated). (F) Representative timelapse fluorescence images of cells expressing Bik1-3mCherry, GFP-Tub1, and Dyn1^MT-U^-3YFP (arrowheads, instances of plus ends with Bik1 foci statically associated with the cortex for 27-28 minutes). Cartoons represent cell in first frame of movie (aMT, astral microtubule; SPB, spindle pole body; magenta circles, plus end Bik1 foci statically associated with cortex). See Video S1 for additional examples. (G and H) Plots depicting frequency (G; for events ≥ 3 frames) and duration (H) of plus end-cortex encounters in cells with indicated dynein and/or Num1 allele. P values were calculated from Z scores (for proportion data) or by using a Mann-Whitney test (for intensity values and microtubule-cortex duration values).

The pattern of Pac1 localization in each of these yeast strains reflects that of the wild-type and mutant dyneins (Fig. 1A-C, hatched bars). Of note, whereas neither wild-type nor *dyn1*^*MT-B*^ cells possess Pac1 foci at the cell cortex, a large fraction of *dyn1*^*MT-U*^ cells do, almost all of which colocalize with dynein^MT-U^-3YFP (Fig. 1A and C). Given that Pac1 is never observed at the cortex in wild-type cells, we wondered whether Bik1 (homolog of human CLIP-170) also localizes to cortical sites in *dyn1*^*MT-U*^ cells. Bik1 is required for plus end association of dynein and Pac1, and likely makes a tripartite complex with these proteins at plus ends^19,24,27^. Imaging *dyn1*^*MT-U*^ cells expressing Bik1-3mCherry revealed that this protein also ectopically localizes to cortical sites (Fig. 1D). In fact, whereas some cortical Bik1 foci were not associated with microtubules (see magenta arrowheads and bars in Fig. 1D and E), others were simultaneously associated with the cortex and a microtubule plus end (see red arrowheads and bars in Fig. 1D and E). Time-lapse imaging revealed that plus ends remained anchored at cortical sites in *dyn1*^*MT-U*^ cells for a mean duration of 13.7 minutes (compared to 2.3 and 2.7 min for wild-type and *dyn1*^*MT-B*^ cells, respectively), with some lasting throughout the entire imaging period (30 minutes; Fig. 1F-H, and Video S1). We confirmed the plus ends were anchored via canonical cortical dynein complexes by deleting Num1, which resulted in a large reduction in these events (Fig. 1G and H).

Taken together, these data indicate that dynein is in a microtubule-unbound conformation at plus ends, and that it must switch to a microtubule-bound state to dissociate from Pac1. Failure to do so results in dynein remaining bound to Pac1, Bik1, and the microtubule plus end.

### Allostery within the dynein motor domain accounts for differential Pac1 affinity

To determine the minimal region of dynein that is sufficient to exhibit this differential Pac1 affinity, we assessed the localization of a dynein motor domain truncation (Fig. 2A). We previously found that a region encompassing the AAA+ ring and most of the linker element – dynein_MOTOR_ – is sufficient for Pac1 binding, and thus for localizing to plus ends in cells^37^. This dynein fragment is missing the very N-terminal region of the linker that was previously found to encounter Pac1 during its powerstroke (see arrow, Fig. 2A, right)^29^. We introduced the SRS_CC_ into dynein_MOTOR_ to generate 3YFP-tagged MT-U and MT-B variants, and assessed the extent of their localization in live cells. Consistent with previous studies, dynein_MOTOR_-3YFP was found at a greater fraction of microtubule plus ends than the full-length dynein complex (compare Fig. 1B to 2C)^37^. We observed an even greater extent of plus end binding for dynein_MOTOR_^MT-U^ -3YFP, while dynein_MOTOR_^MT-B^-3YFP was present in fewer cells, and with a lower fluorescence intensity (Fig. 2B and C). Thus, the structural determinants that account for differential Pac1 affinity are contained within the motor domain.

**Figure 2.**
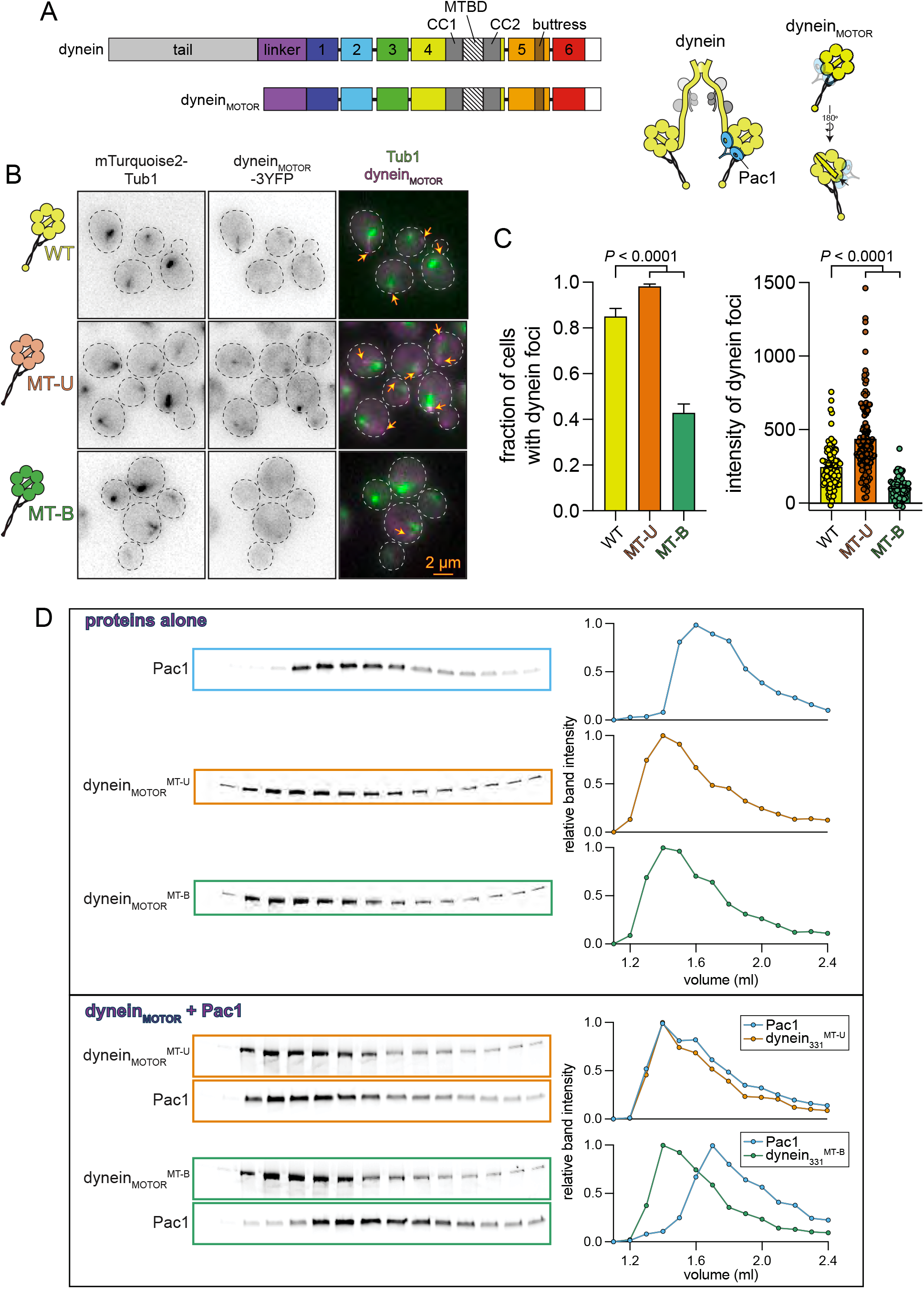
The dynein motor domain is sufficient for microtubule-binding-induced allostery. (A) Schematic and cartoon depictions of full-length and the truncated dynein motor domain used here (CC, coiled-coil; MTBD, microtu-bule-binding domain). Arrow on cartoon indicates truncated dynein linker that does not contact Pac1^29^. Note the truncated motor domain lacks the tail, which is required for Num1 and dynactin binding^37^. (B) Representative fluorescence images of cells expressing mTurquoise2-Tub1 and indicated dynein motor domain fragment (arrows, plus end foci). (C) Plots depicting frequency and intensity of indicated dynein foci, which were scored from timelapse movies. *P* values were calculated from Z scores (for proportion data) or by using a Mann-Whitney test (for intensity values). (D) Analytical size exclusion chromatography analysis showing proteins alone (top), or mixed prior to running on a Superdex 5/150 (bottom). Plots depict band intensity profiles. Gels and analysis are representative of at least 3 independent replicates.

We next asked whether dynein and Pac1 are sufficient to exhibit this behavior *in vitro*. To this end, we purified dynein_MOTOR_ fragments from yeast, mixed them with purified Pac1 in the presence of saturating levels of ATP (3 mM; to roughly approximate *in vivo* conditions), and then applied them to a size exclusion chromatography column. This revealed that Pac1 comigrated with dynein_MOTOR_^MT-U^ to a significantly greater extent than dynein_MOTOR_^MT-B^, indicating that Pac1 binds dynein_MOTOR_^MT-U^ with greater affinity than dynein_MOTOR_^MT-B^ (Fig. 2D).

To validate these findings, we employed mass photometry, a microscopy-based single molecule method that permits determination of the masses of protein species within a mixture^38^. Analysis of each protein alone revealed that the large majority of each had mass values consistent with dimeric Pac1, and monomers of each dynein (Fig. 3A). We then mixed Pac1 with equal concentrations of either wild-type dynein_MOTOR_, dynein_MOTOR_^MT-B^, or dynein_MOTOR_^MT-U^, and assessed the proportion of species that resulted. In the presence of 1 mM ATP, we noted an approximately two-fold greater proportion of dynein_MOTOR_^MT-U^-Pac1 complexes (see ∼520 kDa peak) than dynein_MOTOR_^MT-B^-Pac1 complexes (Fig. 3B and C, “ATP”; also see Fig. 3D). This was also true across a range of Pac1 concentrations (Fig. S2A). Interestingly, this analysis revealed that wild-type dynein_MOTOR_ bound to Pac1 with an affinity that was almost identical to dynein_MOTOR_^MT-U^ (Fig. 3D and S2B), which is consistent with the notion that this mutant mimics wild-type microtubule-unbound dynein.

**Figure 3.**
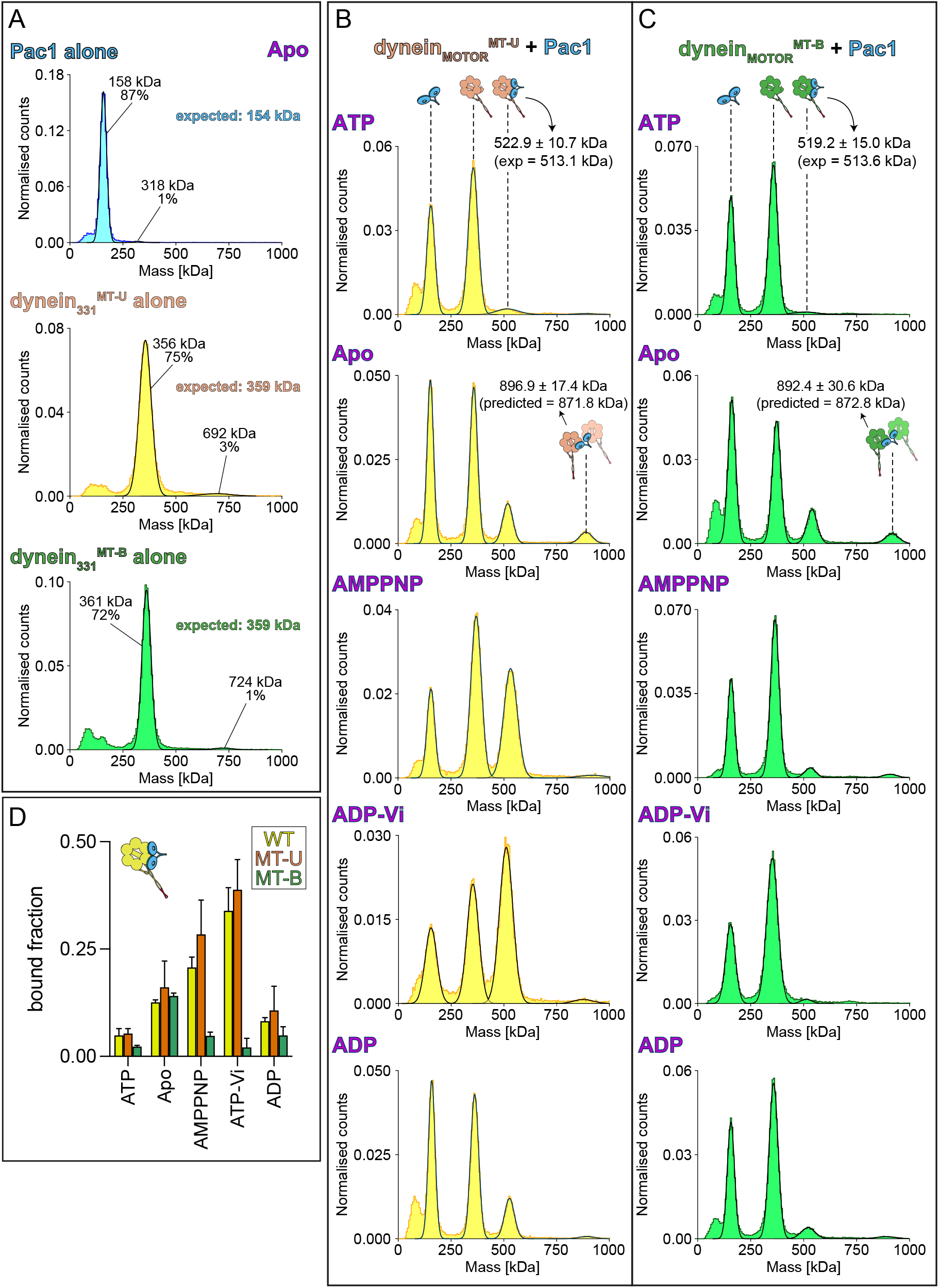
Mass photometric analysis of Pac1-dynein_MOTOR_ binding. (A) Proteins purified from yeast were diluted into assay buffer without nucleotide, and movies were acquired on a Refeyn TwoMP immediately thereafter. The masses of protein species landing on the glass coverslip were empirically determined by converting particle contrast to mass following a calibration routine in the Refeyn software. Fitting of raw data, which identifies mean mass values for each species, and relative fraction of particles with indicated mass, was performed in Discover MP. Note the majority of Pac1 exists as a dimer, while the motor domains are largely monomeric. (B and C) Histograms depicting relative fraction of Pac1 alone, dynein_MOTOR_ alone (MT-U or MT-B), or dynein_MOTOR_-Pac1 complex. Equimolar concentrations of Pac1 and either dynein_MOTOR_^MT-U^ (B) or dynein_MOTOR_^MT-B^ (C; 25 nM each) were mixed in assay buffer with indicated nucleotide (1 mM), incubated for 1-2 minutes, and then diluted to 5 nM on a coverslip mounted on a Refeyn TwoMP. Movies were acquired immediately thereafter, and mass analysis was performed using Discover MP. Note those particles within the 523 kDa peak correspond to 1 Pac1 dimer:1 dynein_MOTOR_ monomer complexes, while those within the 897 kDa peak likely correspond to 1 Pac1 dimer:2 dynein_MOTOR_ complexes (see cartoon schematic above each peak). Plots depict representative data of at least 3 independent replicates for each. (D) The relative fraction of 1 Pac1 dimer:1 dynein_MOTOR_ monomer complexes are plotted (mean ± SD). See Figure S2B for representative mass histograms with the wild-type dynein_MOTOR_ protein with and without Pac1.

### The nucleotide-bound state of dynein affects its affinity for Pac1

Throughout its mechanochemical cycle, the dynein motor domain undergoes a series of conformational changes that are largely a consequence of the bound nucleotides^39,40^. Previous studies found that dynein-LIS1 binding is enhanced by treatment with ATP and vanadate^31^, which results in an ADP-Pi-like state^41^. We wondered how different nucleotides might alter the ability of either MT-U or MT-B to bind Pac1. To this end, we repeated our mass photometry experiments with either no nucleotide (“apo”), AMPPNP (non-hydrolyzable ATP), ATP + vanadate (Vi), or ADP. As expected, ADP-Vi indeed enhances Pac1 binding for both wild-type dynein_MOTOR_ and dynein_MOTOR_^MT-U^ (Figure 3B-D and S2B). However, Pac1-dynein_MOTOR_^MT-B^binding is unaffected by ADP-Vi. AMPPNP also strongly enhances Pac1 binding to dynein_MOTOR_ and dynein_MOTOR_^MT-U^, but only has a minor effect on dynein_MOTOR_^MT-B^. Interestingly, apo conditions led to a situation in which all three dynein_MOTOR_ fragments bound to Pac1 with similar affinities. Finally, ADP had a minor enhancing effect for Pac1 binding to all three fragments. Given the similar response of dynein_MOTOR_ and dynein_MOTOR_^MT-U^ to Pac1 binding for all conditions, these data further support the notion that these two fragments are structurally and biochemically similar, whereas the dynein_MOTOR_^MT-B^ mutant is distinct, and may represent the bona fide microtubule-bound state of dynein.

We noted that a unique ∼890 kDa protein species was apparent in our mass photometry data in a nucleotide-dependent manner. This species was apparent for the wild-type motor (Fig. S2B), and was equally pronounced for both mutants in apo conditions (Fig. 3B and C). In light of the mass of this species, we hypothesize it represents two dynein monomers bridged by a Pac1 dimer. It is unclear what this complex represents, but suggests that Pac1 has the capacity to link distinct dynein molecules or complexes.

In light of the inability of ADP-Vi to affect the Pac1-binding affinity of dynein_MOTOR_^MT-B^, we wondered whether this mutant can bind Vi. To address this, we mixed the purified dynein_MOTOR_ fragments with ATP ± Vi, and exposed the mixtures to ultraviolet light (Fig. S3A). Although both wild-type and dynein_MOTOR_^MT-U^ underwent Vi-dependent photocleavage indicative of Vi binding to AAA1, the dyneinMOTORMT-B mutant did not, indicating that the microtubule-bound conformation of dynein has a low affinity for Vi (and by extension, Pi; Fig. S3B and C). Thus, treatment with ATP + Vi does not enhance the Pac1-binding affinity of dynein_MOTOR_^MT-B^ because it is unable to bind Vi.

Although dynein possesses 6 AAA domains, only 4 of them are competent for nucleotide binding (AAA1-4), whereas 3 possess hydrolysis activity (AAA1, 3 and 4)^42-46^. Studies indicate that AAA1 is the main site for ATPase activity^42,45,47^, with AAA3 and AAA4 playing regulatory roles during motility and force production^48-51^. Previous studies have also found that the nucleotide state of AAA3 may play a direct role in modulating dynein-Pac1 affinity^52,53^. To determine whether ATP binding or hydrolysis at AAA1, 3 or 4 might affect the ability of dynein_MOTOR_^MT-B^ or dynein_MOTOR_^MT-U^, we mutated either Walker A or Walker B (“WA”, or “WB”; responsible for ATP binding and hydrolysis, respectively) in AAA1, 3 and 4, and assessed the extent of their plus end binding. Preventing ATP binding, but not hydrolysis, at AAA1 reduced plus end binding of the MT-U and MT-B mutants, while preventing ATP binding and hydrolysis at AAA3 reduced plus end binding of both (Fig. S3D and E). This suggests that AAA1 is in an ATP-bound state when dynein is at plus ends, but that this is not sufficient to rescue plus end (and thus Pac1)-binding of the MT-B mutant. These data are consistent with our observations that AMPPNP stimulates dynein-Pac1 binding (Fig. 3D). Given the reduced plus end-binding of the AAA3 WA and WB MT-U and MT-B mutants, these data suggest that AAA3 is in neither an apo nor an ATP state when dynein is at plus ends, but potentially an ADP or ADP-Pi state. Finally, our data suggest that the nucleotide-binding state of AAA4 is relatively inconsequential for dynein-Pac1 binding.

### Microtubule-binding induced allostery governing LIS1 affinity is conserved

We sought to determine whether the phenomenon we have described thus far is conserved with human proteins. To this end, we purified LIS1 and corresponding human dynein_MOTOR_, dynein_MOTOR_^MT-U^, and dynein_MOTOR_^MT-B^fragments from insect cells, and assessed their binding via mass photometry. This revealed a very similar approximately 2-fold difference in LIS1-binding affinity between the two mutants, and similar LIS1-binding affinities for dynein_MOTOR_ and dynein_MOTOR_^MT-U^ (Fig. 4B-D). Repeating the binding experiments in the presence or absence of different nucleotides revealed an almost identical response of the human proteins to LIS1 binding as the yeast proteins: a strong enhancement of binding by AMPPNP and ADP-Vi for the wild-type and dynein_MOTOR_^MT-U^ fragments, but not dynein_MOTOR_^MT-B^ (Fig. 4D and S2C). The only notable differences between the yeast and human proteins were a somewhat stronger enhancement of LIS1 binding for both mutants by the apo state, and a more pronounced stimulation by ADP. In summary, these data indicate that microtubule-binding induced conformational changes also reduce the affinity of human dynein for LIS1.

**Figure 4.**
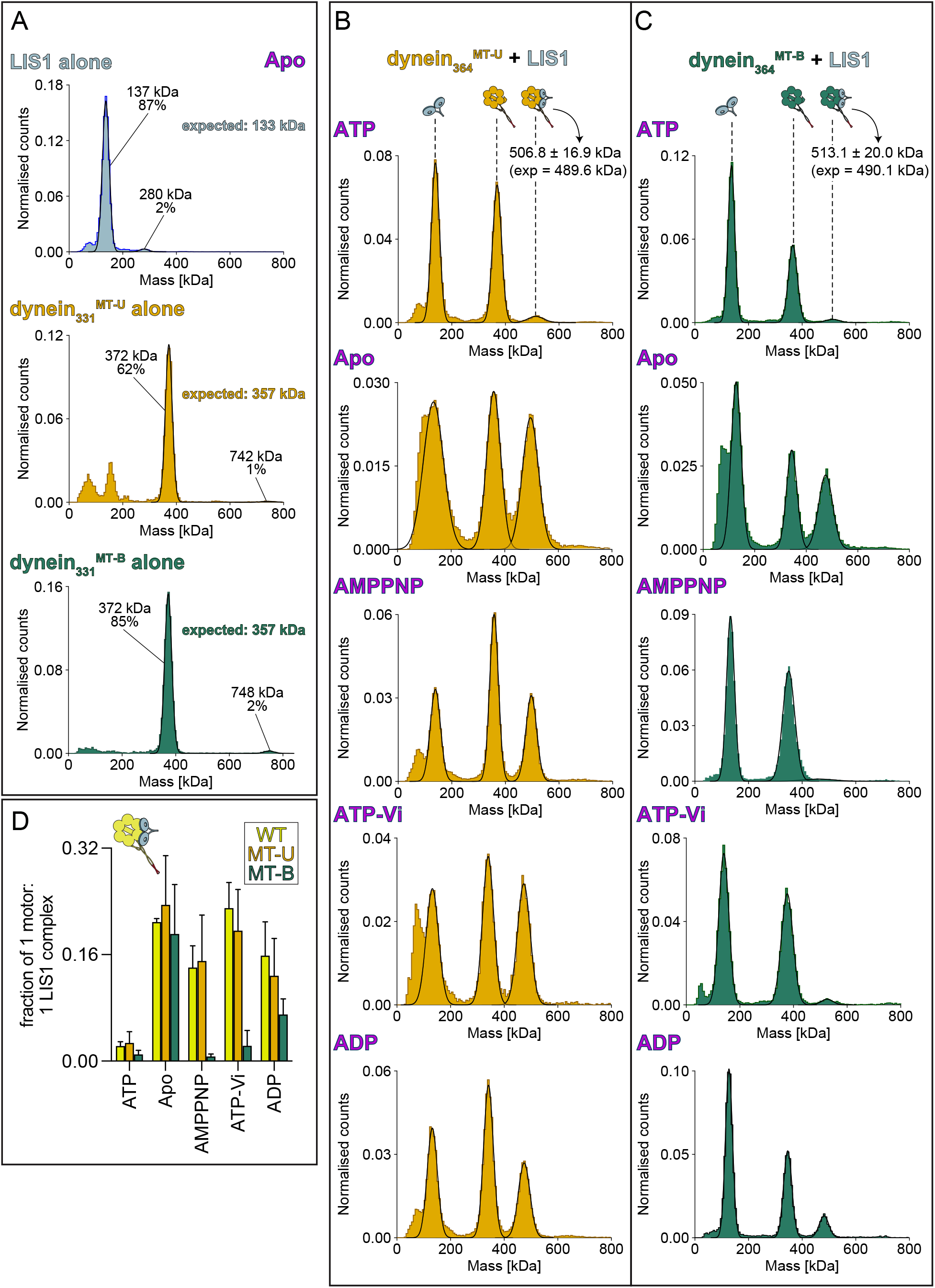
Mass photometric analysis of human LIS1-dynein_MOTOR_ binding. (A) Proteins purified from insect cells were diluted into assay buffer without nucleotide, and movies were acquired on a Refeyn TwoMP immediately thereafter, as described in Figure 3. Note the majority of LIS1 exists as a dimer, while the motor domains are largely monomeric. (B and C) LIS1 and either human dynein _MOTOR_^MT-U^ (B) or dynein _MOTOR_ ^MT-B^ (C) were mixed in assay buffer with indicated nucleotide (to 25 nM each), incubated for 1-2 minutes, and then diluted to 5 nM on the Refeyn TwoMP. Movies were acquired immediately thereafter, and mass analysis was performed using Discover MP. Note those particles within the 507 kDa peak correspond to 1 LIS1 dimer:1 dynein_MOTOR_ monomer complexes (see cartoon schematic above each peak). We did not observe protein species that would correspond to the 1 LIS1 dimer:2 dynein_MOTOR_ complexes observed with yeast proteins in Figure 3. Plots depict representative data of at least 3 independent replicates for each. (D) The relative fraction of 1 LIS1 dimer:1 dynein_MOTOR_ monomer complexes are plotted (mean ± SD). See Figure S2C for representative mass histograms with the wild-type human dynein_MOTOR_ protein with and without LIS1.

### Cryo-EM structures of human MT-B and a LIS1-bound MT-U dynein

Our data thus far indicate that dynein_MOTOR_^MT-U^ behaves almost identically to wild-type dynein with respect to LIS1-binding, but that dynein_MOTOR_^MT-B^ exists in a low LIS1-affinity state. To determine the structural basis for this behavior, we obtained 3.4 and 3.2 Å cryo-EM structures for human dynein_MOTOR_^MT-B^ alone and a LIS1-bound dynein_MOTOR_^MT-U^, respectively (Fig. 5A, S4A-J, and Table S1). While dynein_MOTOR_^MT-B^ was frozen in the presence of ATP, we froze dynein_MOTOR_^MT-U^ in the presence of ATP and Vi to enrich for LIS1-bound complexes. The resolution of our structures permitted us to unambiguously assign nucleotide density to all 4 binding pockets in both the dynein motors (Fig. S5A and E). Of note, although density for ADP-Vi was apparent in AAA1 in the dynein_MOTOR_^MT-U^ structure, the AAA1 pocket of dynein_MOTOR_^MT-B^ was bound to ADP, suggesting that the microtubule-bound state of dynein has a high affinity for ADP at AAA1. This is consistent with a recent report in which ADP was observed at AAA1 for a native microtubule-bound dynein-dynactin-adaptor complex^54^.

**Figure 5.**
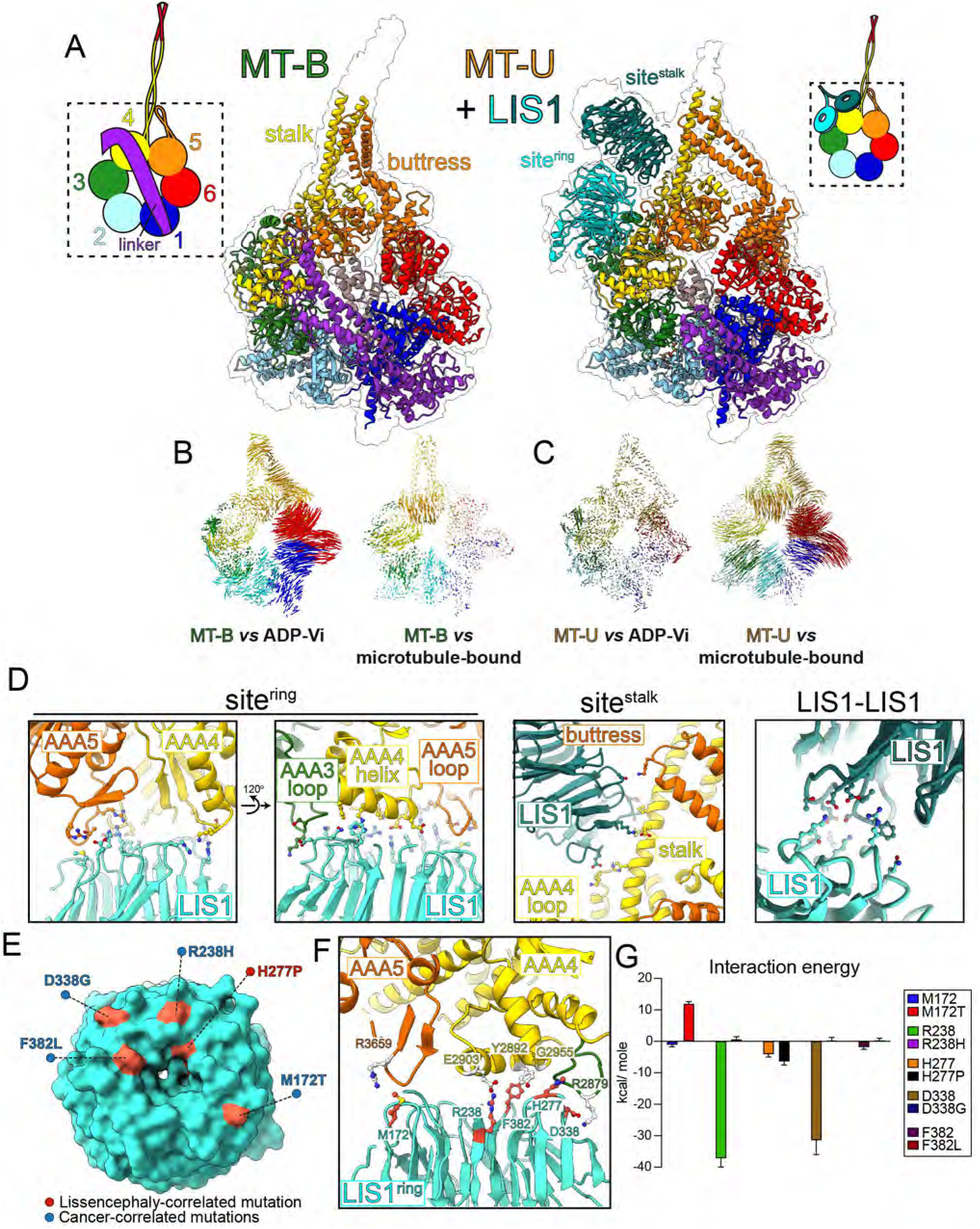
Cryo-EM structures of human dynein_MOTOR_^MT-B^ and a LIS1-bound dynein^MT-U^. (A) Molecular models of dynein _MOTOR_^MT-B^ (solved in the presence of ATP) and LIS1-bound dynein _MOTOR_^MT-U^ (solved in the presence of ATP and Vi) with corresponding density maps (indicated with outlines). Subdomains are color-coded as indicated by cartoon. (B and C) Vector maps depicting pairwise alpha carbon interatomic distances between dynein_MOTOR_^MT-B^ (B) or dynein_MOTOR_^MT-U^ (C) with either the human ADP-Vi-bound dynein-2 crystal structure (4RH7)^41^, or the native microtubule-bound dynein-1 cryo-EM structure (7Z8G)^54^. Structures were globally aligned after removal of the linkers. The length of the lines are proportional to the calculated interatomic distances. Note the strong similarities between dynein_MOTOR_^MT-B^ and the micro-tubule-bound dynein (but not with 4RH7), and that of dynein_MOTOR_^MT-U^ with the ADP-Vi-bound dynein (but not with 7Z8G). (D) Close-up views of the main contact points between LIS1 with sitering and sitestalk (as indicated), and between the two LIS1s within the homodimer. Also see Video S2. Residues with atoms shown are those determined to mediate contacts (see Figure S6B). (E) Surface view showing sitering-bound LIS1 with disease-correlated residues highlighted. (F) Close-up view of contact points between disease-correlated residues on LIS1 and sitering. (G) Results of molecular dynamics simulation depicting energy of interaction between wild-type or mutated residue, as indicated. See Figure S6D for graphical depiction of MD simulation data.

Comparisons to published structures revealed various degrees of differences and similarities (Fig. 5B and C, and S5B and G). Notably, that which most closely resembles dynein_MOTOR_^MT-B^ is the native microtubule-bound dynein described above (Fig. 5B and S5C). In fact, the minor differences between these two structures can be accounted for by the presence of AMPPNP instead of ADP at AAA3 in the native microtubule-bound dynein (Fig. S5D). Consistent with our data indicating that the dynein_MOTOR_^MT-U^ mutant behaves like wild-type dynein in the absence of microtubules, the dynein_MOTOR_^MT-U^ structure very closely resembles that of the ADP-Vi-bound dynein-2 (Fig. 5C). These data indicate that our mutants indeed reflect microtubule-bound and - unbound conformations, and that the microtubule-bound state is distinct from that adopted by a microtubule-unbound dynein in the presence or absence of various nucleotide analogs (Fig. S5B).

Our structure of the dynein_MOTOR_^MT-U^ -LIS1 complex revealed the monomeric dynein motor bound to two LIS1 WD40 beta-propellers (Fig. 5A). Given our mass photometry data indicate a 1:1 binding stoichiometry (1 LIS1 dimer:1 dynein motor domain), these two beta-propellers are likely from the same LIS1 homodimer (see Fig. S6A, right, and S6C for model). Consistent with previous structures of yeast dynein bound to a Pac1 dimer^52,55^, the LIS1 beta-propellers were bound to two sites on the human dynein motor (Fig. 5A): one at the interface of AAA3 and AAA4 (referred to as site^ring^), and the other at the base of the stalk (site^stalk^). Binding of LIS1 to site^ring^ involves a surface exposed helix in AAA4, a short loop within AAA5, and a region of a longer loop within AAA3, while binding at site^stalk^ involves residues along the stalk, part of a long loop within AAA4, and residues at the tip of the buttress (Fig. 5D, S6B, S7 and Video S2). As with the yeast counterparts, we also noted contacts between the two LIS1s, which have been shown to be important for yeast dynein function^55^.

Inspection of the regions of LIS1 that contact site^ring^ and site^stalk^ revealed numerous residues distributed over the face of the two beta-propellers (Fig. S6B, S8 and Video S2). Although one of these residues (H277) has been found to be mutated in a patient with lissencephaly^56^, four others have been found to be mutated in cancer patients (Fig. 5E and F; M172T, R238H, D338G, and F382L)^57^. We used molecular dynamics (MD) simulations to analyze the potential consequences of these mutations on the dynein-LIS1 interaction, and found that the cancer-correlated mutations all decrease the energy of interaction to varying degrees, while the lissencephaly-correlated mutation did not (Fig. 5G and S6D). These data suggest that weakened LIS1-dynein interactions caused by these mutations may be linked to tumorigenesis.

An analysis of our dynein_MOTOR_^MT-U^-LIS1 cryo-EM dataset revealed that a subset of these dyneins were bound to only 1 LIS1 (36%), while the remaining were either bound to 2 (29%) or “1.5” molecules (34%), in which a strong density was apparent for only one of the LIS1 molecules, with a weaker density corresponding to the 2^nd^ LIS1, which is indicative of flexibility of this latter molecule (Fig. S4F). All LIS1-bound dyneins possess clear density at site^ring^, indicating this is the primary binding site, while the presence of the site^stalk^-bound LIS1 was variable, suggesting that this site is the lower affinity LIS1-binding site on dynein. Interestingly, local resolution analysis of the three classes (those bound to 1, 1.5, and 2 LIS1s) revealed that the density for the site^ring^-bound LIS1 is best for the 1.5 and 2 LIS1-bound dyneins (as apparent from the resolution of the bound LIS1; Fig. S9A). Moreover, all three classes exhibit clear density of ADP within the AAA3 binding pocket, suggesting that the nucleotide state is not causative of these differences (Fig. S9B). These observations suggest that the binding of LIS1 to site^stalk^, which appears to be rate-limiting, stabilizes the entire LIS1 dimer-dynein complex. In addition to the improved resolution of LIS1, we also noted that two regions within dynein at site^stalk^ also exhibit greater resolution when more than 1 LIS1 is present: the AAA4 loop (residues 3111-3138), and the tip of the buttress, suggesting these regions become less flexible when bound to LIS1 (Fig. S9B; also see below).

### Cryo-EM structure of human MT-U dynein without LIS1

The reduced flexibility of the LIS1-binding regions of dynein (Fig. S9B), as well as previously published work suggest that LIS1 binding may affect the conformation of dynein^30,52^. It remains controversial whether this binding affects dynein’s mechanochemistry and/or microtubule-binding behavior^9^. We reasoned that if LIS1 were to impact the biochemical behavior of dynein, its binding would cause structural changes reflective of these activities. To determine if this is the case, we solved a 2.9 Å cryo-EM structure of dynein_MOTOR_^MT-U^ in the absence of LIS1, but in the presence of ATP and Vi (to allow an accurate comparison with the LIS1-bound dynein). With a few exceptions, this revealed a structure that was almost identical to the LIS1-bound dynein_MOTOR_^MT-U^ (Fig. 6A and B). Notably, the conformation of the nucleotide binding pocket at AAA3, which was clearly bound to ADP, appears unchanged between the LIS1-bound and unbound dyneins (Fig. 6C). Among the exceptions are the following small differences at the LIS1-binding sites (Fig. 6D): the tip of the buttress is shifted 3.1 Å toward the site^stalk^-bound LIS1, which results in a 1.7 Å shift of CC2 toward CC1 of the stalk; and, the AAA5 loop is shifted 1.4 Å away from the site^ring^-bound LIS1. The overall conformational similarities between these two dynein structures are consistent with recent findings that LIS1 does not in fact impact dynein’s mechanochemistry or microtubule-binding behavior^8^. Rather, our findings indicate that dynein’s conformational state impacts its ability to bind LIS1, but not vice versa.

**Figure 6.**
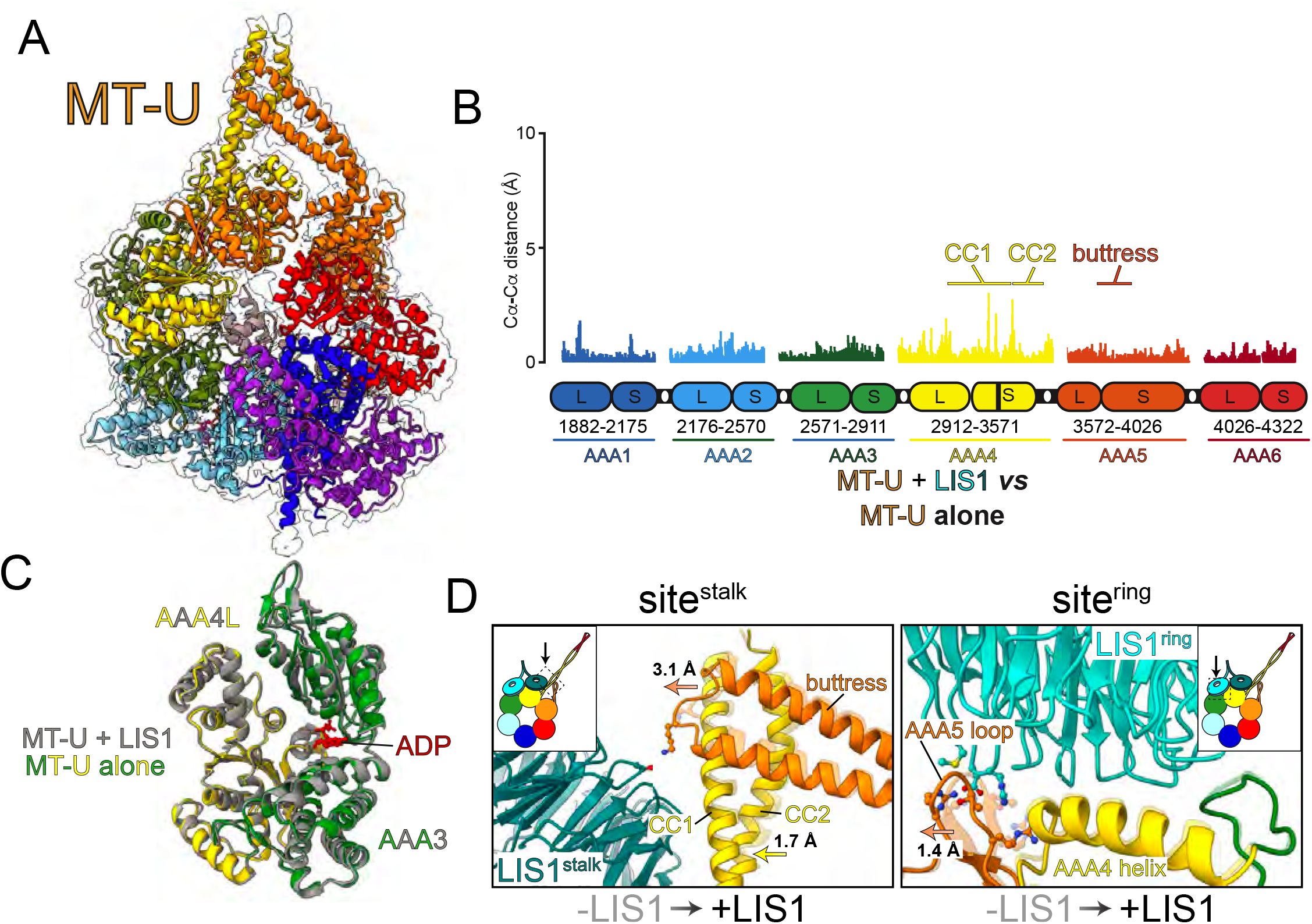
Cryo-EM structure of human dynein_MOTOR_^MT-U^ alone. (A) Molecular model of dynein_MOTOR_^MT-U^ (solved in the presence of ATP and Vi) with corresponding density map (indicated with outline). Subdomains are color-coded as indicated by cartoon shown in Figure 5A. (B) Plot depicting pairwise alpha carbon interatomic distances between the dynein_MOTOR_ ^MT-U^ with and without LIS1. Note the high degree of similarity between the two structures, with minor exceptions in CC1 and CC2 (see text). (C) AAA3-AAA4L domains from dynein_MOTOR_ ^MT-U^ with (grey) and without LIS1 (green and yellow) overlaid to depict the high degree of structural similarity. (D) Close-up views illustrating the differences in dynein structure with and without LIS1 at the contact points between dynein and LIS1. The structure without LIS1 is depicted with reduced opacity compared to that with LIS1. Note the small shifts in the buttress tip toward sitestalk-bound LIS1, and of the AAA5 loop away from sitering-bound LIS1.

### Microtubule-binding induced changes at the ring site account for reduced LIS1 affinity

Global alignment of MT-U and MT-B reveal the changes initiated by microtubule binding and the consequent CC1/CC2 helix sliding in the stalk^58^. Movement of CC2 with respect to CC1 causes the tip of the buttress to shift away from the AAA+ ring. This leads to a deep kink in the middle of the buttress, and a consequent rigid body movement of the AAA5S-AAA6L subdomains. This causes the AAA+ ring to adopt a more ‘open’ state that can no longer coordinate Pi binding at AAA1 (Video S3 and Figure S5H). We wondered whether these structural changes spanning one side of the AAA+ ring (AAA1, AAA5, AAA6) lead to allostery on the other side (*i*.*e*., at site^ring^ and site^stalk^) that would influence dynein-LIS1 affinity. Local alignment of MT-U and MT-B using AAA4-AAA5 revealed several notable changes at both LIS1 binding interfaces, including the following: at site^stalk^, the tip of the buttress is shifted 10.3 Å away from LIS1 (Fig. 7A); at site^ring^, both the AAA4 helix (residues 2886-2903) and the AAA3 loop (residues 2875-2880) are shifted 4.4-6.5 Å away from the AAA5 loop (residues 3654-3661), thus increasing the spacing between these three elements that all make direct contacts with LIS1 (Fig. 5D and 7B; also see Video S4). This latter change is likely sufficient to significantly weaken the binding affinity of LIS1 to site^ring^.

**Figure 7.**
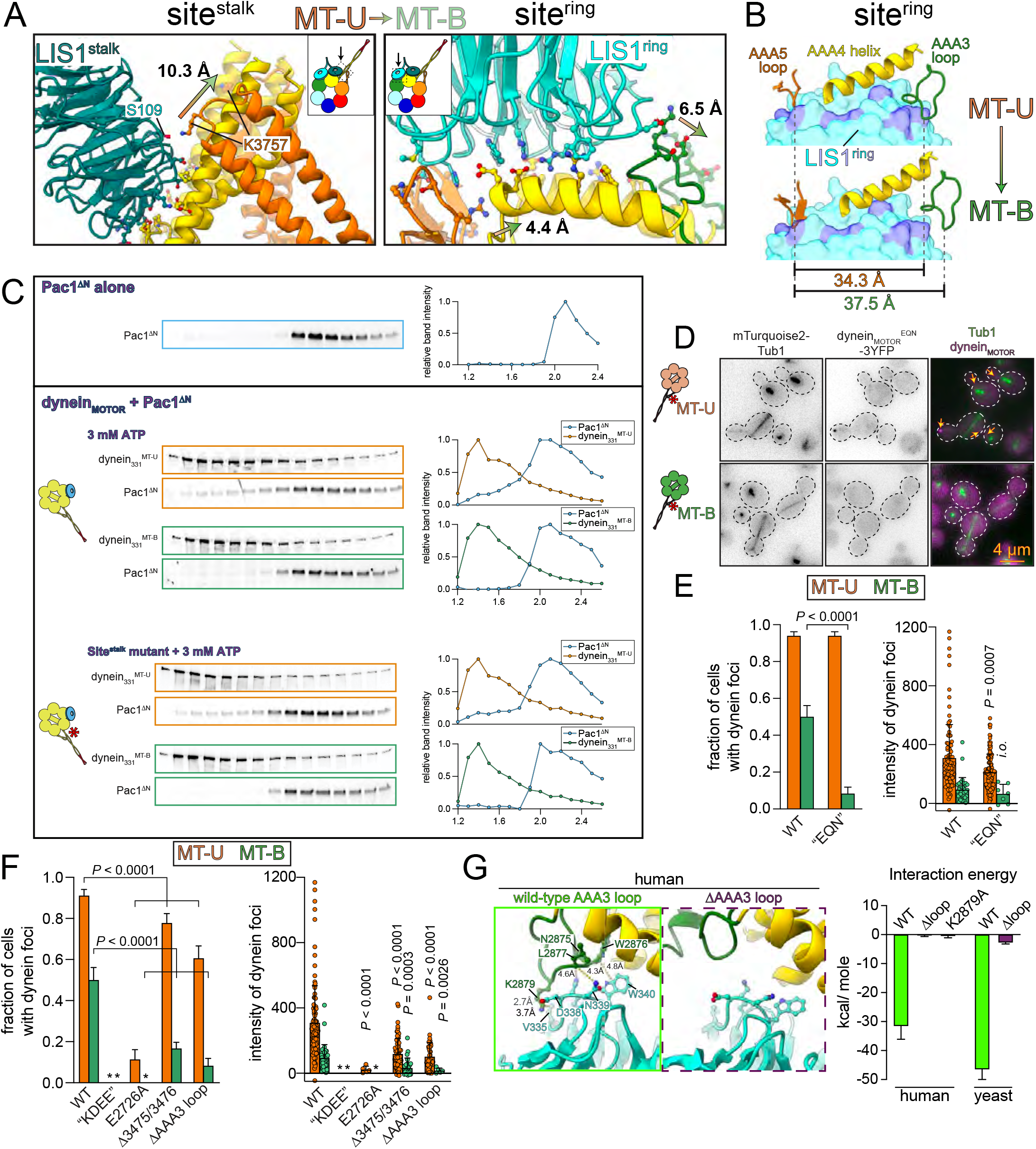
Changes at site_ring_ account for reduced LIS1/Pac1 binding affinity. (A) Close-up views illustrating the differences in structure between dynein_MOTOR_^MT-U^ and dynein_MOTOR_^MT-B^ at sitestalk and sitering. The dynein_MOTOR_^MT-B^ struc-ture is depicted with reduced opacity compared to dynein_MOTOR_^MT-U^. Notable differences are indicated with arrows with values indicating interatomic distances between relevant residue sidechains. Residues with atoms shown are those determined to mediate contacts between dynein and LIS1. (B) Summary of major changes at sitering. Residues on LIS1 space-filling model shown in purple are those that make contact with sitering. Note the increased distance between the AAA5 loop and the AAA4 helix and AAA3 loop (that comprise critical contact points with LIS1) that results in an approximate 3.2 Å distortion of the binding site. (C) Analytical size exclusion chromatography analysis showing monomeric Pac1^∆N^ alone (top), or mixed with indicated yeast dynein_MOTOR_ (either wild-type version of MT-U/MT-B, middle, or those with “EQN” mutations, bottom) prior to running on a Superdex 5/150 (bottom). Plots depict band intensity profiles. Gels and analysis are representative of at least 3 independent replicates. Note the apparent binding between Pac1^∆N^ and dynein_MOTOR_^MT-B^, but not with dynein_MOTOR_^MT-B^. (D) Representative fluorescence images of cells expressing mTurquoise2-Tub1 and indicated dynein motor domain fragment with “EQN” mutations (arrows, plus end foci). (E and F) Plots depicting frequency and intensity of indicated dynein foci, which were scored from timelapse movies (*i*. *o*, in. sufficient observations; *, no foci observed). (G) Results of molecular dynamics simulation depicting energy of interaction between wild-type or mutant human or yeast dynein with LIS1 or Pac1.

Our data suggest that LIS1-binding to dynein is initiated at site^ring^, and followed by site^stalk^ (see above and Fig. S9). This is further supported by the fact that a monomeric Pac1 binds predominantly at site^ring^ with no apparent binding at site^stalk29^. Thus, we sought to determine if structural changes at site^ring^ are responsible for the altered Pac1 and LIS1-binding affinity. To this end, we specifically interrogated this site by assessing the binding between dynein and a monomeric Pac1 mutant (lacking its N-terminal dimerization domain; Pac1^∆N^). We found that Pac1^∆N^ comigrates with dynein_MOTOR_^MT-U^ via size exclusion chromatography to a significantly greater extent than dynein_MOTOR_^MT-B^, indicating that the Pac1 monomer indeed exhibits differential binding affinities for these two mutants, much like the Pac1 dimer (Figure 7C). To ensure that Pac1^∆N^ was binding to site^ring^ we repeated the binding assay using dynein variants with three point mutations that interfere with Pac1-site^stalk^ binding: E3012A, Q3014A, and N3018A (“EQN” mutant)^52^. This revealed an identical binding disparity of Pac1 for the MT-U and MT-B mutants (Fig. 7C, bottom). We validated these findings in cells by assessing the localization of EQN dynein_MOTOR_^MT-U^-3YFP and dynein_MOTOR_^MT-B^-3YFP mutants to microtubule plus ends. Although both had reduced plus end localization with respect to the wild-type versions of each, the two proteins exhibited localization disparities even more extreme than the wild-type MT-U and MT-B (1.8X difference in frequency for wild-type, versus 11.7X for the EQN mutants; Fig. 7D and E). These data are consistent with a role for site^stalk^ in dynein-Pac1 binding, and indicate that site^ring^ is indeed undergoing a conformational change that weakens its affinity for Pac1 and LIS1 upon microtubule binding.

We next focused on the structural elements at site^ring^ that may account for the microtubule-binding induced Pac1/LIS1-dissociation: the AAA4 helix, the AAA3 loop, and the AAA5 loop, which move with respect to each other upon microtubule-binding (Fig. 7A and B, and Video S4). Consistent with the importance of the AAA4 helix in dynein-Pac1 binding, mutating either four residues (K2721A, D2725G, E2726S, and E2727G; referred to as the “KDEE” mutant)^30^ or only one (E2726A; equivalent to E2903 in human dynein) severely reduces plus end binding of both dynein_MOTOR_^MT-U^ and dynein_MOTOR_^MT-B^ in cells (Fig. 7F). Furthermore, deleting two key residues in the AAA5 loop (N3475 and R3476) significantly reduces plus end binding of both MT-U and MT-B dyneins (Fig. 7F)^55^, demonstrating the importance of this surface in Pac1-binding. We next focused on the AAA3 loop as a potential Pac1/LIS1-binding surface that changes in response to microtubule-binding. In support of the importance of this interface, which includes a salt bridge between LIS1 D388 and dynein K2879, MD simulation data reveal that the cancer-correlated D338G mutation in LIS1 significantly reduces binding energy (Fig. 5E-G and S6D). Additional MD simulations reveal that either deleting this loop (∆2875-2880) or mutating K2879 to an alanine in human dynein, or deleting this loop in yeast dynein (∆2678-2703) significantly reduces binding energy between dynein and LIS1/Pac1 (Fig. 7G). Finally, yeast cells expressing dynein_MOTOR_^MT-U^-3YFP and dynein_MOTOR_^MT-B^-3YFP mutants with this loop deleted (∆2698-2703) exhibit significant reductions in the intensities and frequencies of foci (Fig. 7F). These data demonstrate the importance of this loop in the dynein-LIS1 contact, and indicate that the conformational changes at site^ring^ that result from microtubule binding likely account for disruption of the dynein-LIS1 complex. Although our experiments do not exclude the involvement of site^stalk^, they indicate that site^ring^ is at least partially responsible for the microtubule-binding induced change.

## DISCUSSION

Our work reveals insight into the final step of the LIS1-mediated dynein activation pathway, and the consequences of preventing it. We find that microtubule binding by dynein triggers its dissociation from LIS1, and that this is required to uncouple the dynein transport complex from the plus end-targeting machinery (*e*.*g*., Bik1/CLIP-170 in yeast, and EB1 in metazoa)^19,20,25,59^. This dissociation is required for dynein to switch from a plus end-associated state, in which it is only indirectly associated with the microtubule, to a motile state, in which it is directly engaged with the microtubule lattice. Dynein is in a microtubule-unbound conformation when it is associated with the plus end-binding machinery. Preventing its switching to a microtubule-bound state locks dynein in this plus end-associated state in cells. In light of the similar affinity of wild-type and dynein^MT-U^ for Pac1 and LIS1 *in vitro*, it is the microtubule-bound state that is a unique conformational state that exhibits low affinity for Pac1 and LIS1. In addition to revealing the structural basis for the weakened affinity, we find that LIS1 binding to dynein is governed by the conformational state of dynein, but that LIS1 binding has no significant effect on dynein’s conformation, consistent with recent work from our lab showing that Pac1 does not in fact affect dynein mechanochemistry^8^.

Based on our new findings, as well as those from other groups, we posit the following model for Pac1 function in budding yeast (see Video S5): (1) dynein exists predominantly in an autoinhibited phi conformation in the cytoplasm, and stochastically switches to an open state^7,8^; (2) once in an open state, Pac1 binds to dynein due to the increased accessibility of Pac1-binding surfaces, thus preventing dynein from switching back to the phi particle^8,12^; (3) the dynein-Pac1 complex binds to plus end-bound Bik1 in a manner that does not require the dynein MTBD^26^; (4) dynactin associates with open, plus end-bound dynein; (5) the dynein-dynactin complex binds to cortical Num1 receptors, which triggers dynein-microtubule binding, potentially by arranging the motor domains in a parallel configuration^7^; (6) microtubule-binding by dynein triggers a cascade of conformational changes, including a distortion of site^ring^ that weakens its affinity for Pac1; (7) Pac1 dissociates from dynein, thus breaking dynein’s indirect connection to the plus end; (8) dynein-dynactin directly engage with the microtubule, and translocate the mitotic spindle toward the bud neck, the site of cytokinesis in budding yeast. In light of the similarities between the yeast and metazoan systems^9^, and our data with human dynein and LIS1, we posit that a very similar mechanism is at play in animal cells.

Given that dynein is in a microtubule-unbound conformation at plus ends, and that the dynein MTBD is dispensable for this association^26^, our work indicates that Pac1 (and likely LIS1) does not in fact promote plus end binding by impacting dynein’s microtubule-binding behavior. In fact, we show that once dynein makes direct contact with the microtubule, this leads to a consequent dissociation of dynein from Pac1. This likely explains the lack of Pac1/LIS1-dynein colocalization at sites of dynein activity (*e*.*g*., the cell cortex in budding yeast, retrograde-moving endosomes in filamentous fungi)^22-24^. These data are further supported by our cryo-EM structures for dynein^MT-U^ in the absence and presence of LIS1, which show very little differences between them. Thus, LIS1-binding does not appear to impact dynein conformation.

Our work also reveals the first high resolution structure of a human dynein-LIS1 complex, thus highlighting the precise residues that link these molecules together. Our 3D classification analysis of the different LIS1-bound dynein species (*i*.*e*., 1, “1.5”, and 2 LIS1s) reveal insight into the importance of the site^stalk^-bound LIS1: as occupancy of site^stalk^ by LIS1 increases, so does the resolution of the site^ring^-bound LIS1, indicating that this latter contact site is stabilized by the former. These data support an avidity model, in which having two bindings sites on dynein improves LIS1 binding. This is consistent with data indicating that a Pac1 monomer in yeast can rescue function only if overexpressed^30^. The importance of LIS1-LIS1 binding is further highlighted by the presence of disease-correlated missense mutations in the LIS1 LisH dimerization domain (*e*.*g*., F31S, L43S, W55M)^60-62^.

Although our structural analysis reveals the likely basis for dynein-Pac1/LIS1 dissociation, we cannot completely discount other changes that might contribute to this process. For example, a previous study found that the N-terminal region of the dynein linker element encounters Pac1 during its powerstroke^29^. One potential hypothesis is that the linker swing may thus partly account for the dissociation (*i*.*e*., by knocking it off). However, in contrast to this possibility, we find a dynein with a shortened linker that does not encounter Pac1^29^ is sufficient for the dissociation. Furthermore, a previous study found that treatment with AMPPNP leads to a straight post-powerstroke linker^63^. Given our findings that AMPPNP stimulates dynein-Pac1/LIS1 binding, the linker position is likely inconsequential to dynein-Pac1/LIS1 binding. Finally, we cannot rule out changes at site^stalk^ being at least partly responsible for this phenomenon. In fact, given the apparent contact between the tip of the buttress of dynein^MT-U^ and the site^stalk^-bound LIS1, the affinity of LIS1 for this site is likely also weakened by microtubule-binding.

One of our more surprising findings was the inability of dynein^MT-B^ to bind to vanadate. This observation seems to conflict with our ATPase assays indicating this mutant hydrolyzes ATP at a similarly high rate to wild-type dynein in the presence of microtubules. Although it is well-established that microtubule binding by dynein accelerates ATP hydrolysis of the AAA1-bound nucleotide, it is unclear how this is achieved. Our data indicate that microtubule-binding specifically stimulates phosphate release by ‘opening’ the AAA1-AAA2L interface, resulting in the AAA2L arginine finger, which stabilize the Vi, moving 8.5 Å away from the AAA1L Walker B glutamate in the dynein^MT-B^ state (Figure S5H; also see Video S4). It is also interesting to note that dynein^MT-B^ exhibits clear density for ADP at this site, indicating a majority of particles are bound to ADP at AAA1. A recent cryo-EM structure of a native microtubule-bound dynein also identified ADP at this site^54^. The authors attributed this to the presence of AMPPNP at AAA3; however, our near-atomic structures obtained without AMPPNP indicate this is more likely a consequence of AAA1 having a high affinity for ADP in the microtubule-bound state, or that ADP release is a rate-limiting step for microtubule-bound dynein. Future studies will be required to more carefully dissect the role of microtubule-binding and unbinding in dictating the ATP hydrolysis cycle.

## ACKNOWLEDGEMENTS

We are very grateful to Janelia Research Campus for providing fluorescent Halo and SNAP dyes, Andrew Carter and Sami Chabaan for valuable discussions (and for sharing data that was unpublished at the time), and to members of the Markus and DeLuca laboratories for valuable discussions. This work was funded by the NIH/NIGMS (R35GM139483 to S.M.M., and R35GM142959 to K.Z.). We would like to thank Kaifeng Zhou, Jianfeng Lin for the help with cryo-EM data collection at Yale ScienceHill-Cryo-EM facility. The Yale Cryo-EM Resource is funded in part by the NIH grant S10OD023603. We thank Liguo Wang, Jake Kaminsky, Guobin Hu at the Laboratory for BioMolecular Structure (LBMS) for help on cryo-EM data collection. The LBMS is supported by the DOE Office of Biological and Environmental Research (KP1607011). Models have been deposited in the PDB as follows: **XXX, YYY, ZZZ**. The residues labeled in the figures are based on the Cytoplasmic Dynein-1 Heavy Chain 1 sequence (Q14204).

## METHODS

### Media and strain construction

Strains are derived from either W303 (for protein purification) or YEF473 (for cell imaging)^64^. We transformed yeast strains using the lithium acetate method^65^. Strains carrying mutations, insertions (*e*.*g*., SRS^CC^), or tagged components were constructed by PCR product-mediated transformation^66^, by transforming plasmids with recombination or expression cassettes^8,37^, or by mating followed by tetrad dissection. Proper tagging and mutagenesis was confirmed by PCR, and in some cases sequencing. Fluorescent tubulin-expressing yeast strains were generated using plasmids and strategies described previously^67^. Yeast synthetic defined (SD) media was obtained from Sunrise Science Products (San Diego, CA).

### Plasmid and BACmid construction

To construct yeast strains expressing SRS^CC^-containing yeast dynein motor domain fragments (dynein_MOTOR_^MT-U^ and dynein_MOTOR_^MT-B^), we engineered pSM01:Dyn1_MOTOR_-3YFP using Gibson assembly^37,68^. The seryl tRNA sythetase coiled-coil (residue 30-96) was amplified from *T. thermophilus* (strain BB8) genomic DNA and assembled into pSM01:Dyn1_MOTOR_-3YFP to achieve the sequences depicted in Figure S1B (note the presence of 4 additional amino acids in the MT-B mutant with respect to MT-U). All mutants were engineered into this plasmid (*e*.*g*., EQN, KDEE, *etc*), which was digested (with BsaBI/BsiWI; to release the respective dynein open reading frame along with a *TRP1* marker, all of which is flanked with homology arms) and subsequently transformed into *dyn1∆::HIS3* yeast. The entire cassette is integrated into the *DYN1* locus (resulting in HIS^−^/TRP^+^ cells), and is the only source of dynein in these cells. Genomic integration was confirmed by growth on selective solid media (one lacking histidine, and another lacking tryptophan), and by PCR.

The human dynein motor domain (residues 1458-4646) was amplified from pbiG1a:6XHis-ZZ-TEV-SNAPf-DHC (codon optimized for insect cells; a gift from A. Carter)^69^ and assembled into pFastBac:8His-ZZ-TEV-LIS1-SNAPf (replacing LIS1-SNAPf), generating pFastBac:8His-ZZ-TEV-DHC_MOTOR_. The SRS^CC^ from *T. thermophilus* was engineered into this plasmid to generate the MT-U and MT-B mutants. These plasmids (along with pFastBac:8His-ZZ-TEV-LIS1-SNAPf) were individually transformed into DH10 EMBacY cells (Geneva Biotech, Geneva, Switzerland) according to the manufacturer’s protocol. Proper transposition and BACmid generation was confirmed by blue/white screening, and by PCR.

### Protein purification

Purification of yeast dynein (wild-type and mutants; ZZ-TEV-dynein_331_-HALO, or ZZ-TEV-6His-GFP-3HA-GST-dynein_331_-HALO, all under the control of the galactose-inducible promoter, *GAL1p*) or Pac1-SNAP was performed as previously described^8,30,70^. Briefly, yeast cultures were grown in YPA supplemented with 2% galactose, harvested, washed with cold water, and then resuspended in a small volume of water. The resuspended cell pellet was drop frozen into liquid nitrogen and then lysed in a coffee grinder (Hamilton Beach). After lysis, 0.25 volume of 4X lysis buffer (1X buffer: 30 mM HEPES, pH 7.2, 50 mM potassium acetate, 2 mM magnesium acetate, 0.2 mM EGTA) supplemented with 1 mM DTT, 0.1 mM Mg-ATP, 0.5 mM Pefabloc SC (concentrations for 1X buffer) was added, and the lysate was clarified at 310,000 x g for 1 hour. The supernatant was then bound to IgG sepharose 6 fast flow resin (Cytinva) for 2-4 hours at 4°C, which was subsequently washed three times in 5 ml lysis buffer, and twice in TEV buffer (50 mM Tris, pH 8.0, 150 mM potassium acetate, 2 mM magnesium acetate, 1 mM EGTA, 0.005% Triton X-100, 10% glycerol, 1 mM DTT, 0.1 mM Mg-ATP, 0.5 mM Pefabloc SC). To fluorescently label the proteins, the bead-bound protein was incubated with either 10 µM JFX646-HaloTag (for the motors) or JF646-SNAP-tag ligand (for Pac1; Janelia Research Campus) for 30-60 minutes at 4°C. The resin was then washed four more times in TEV digest buffer, then incubated in TEV buffer supplemented with TEV protease overnight at 4°C. Following TEV digest, the protein-containing supernatant was collected using a spin filtration device, aliquoted, flash frozen in liquid nitrogen, and stored at -80ºC.

Human proteins (LIS1-SNAP, or motor domains) were expressed and purified from insect cells (ExpiSf9 cells; Life Technologies) as previously described with minor modifications^3,7,20,69^. Briefly, 4 ml of ExpiSf9 cells at 2.5 × 10^6^ cells/ml, which were maintained in ExpiSf CD Medium (Life Technologies), were transfected with 1-9 µg of bacmid DNA using ExpiFectamine (Life Technologies) according to the manufacturer’s instructions. 4-8 days following transfection, the cells were pelleted, and 1-2 ml of the resulting supernatant (P1) was used to infect ∼150 ml of ExpiSf9 cells (5 × 10^6^ cells/ml). Approximately 65 hours later, the cells were harvested (2000 x g, 20 min), washed with human dynein lysis buffer (50 mM HEPES, pH 7.4, 100 mM NaCl, 10% glycerol, 1 mM DTT, 0.1 mM Mg-ATP, 1 mM PMSF; note ATP was omitted for LIS1 purification), pelleted again (1810 x g, 20 min), and resuspended in an equal volume of same. The resulting cell suspension was drop frozen in liquid nitrogen and stored at -80ºC. For protein purification, 30 ml of additional human dynein lysis buffer supplemented with cOmplete protease inhibitor cocktail (Roche) was added to the frozen cell pellet, which was then rapidly thawed in a 37ºC water bath prior to incubation on ice. Cells were lysed in a dounce-type tissue grinder (Wheaton) using 50-60 strokes. Subsequent to clarification at 310,000 x g for 1 hour, the supernatant was applied to 2 ml of IgG sepharose fast flow resin (GE) pre-equilibrated in human dynein lysis buffer, and incubated at 4ºC for 3-5 hours. Beads were then washed 3 times with with 5-10 ml of human dynein lysis buffer, and 2 times with 5-10 ml of human dynein TEV buffer (50 mM Tris pH 7.4, 150 mM potassium acetate, 2 mM magnesium acetate, 1 mM EGTA, 10% glycerol, 1 mM DTT, 0.1 mM Mg-ATP; note ATP was omitted for LIS1 purification). The beads were incubated with TEV protease overnight at 4ºC. The next morning, the recovered supernatant was collected, concentrated, aliquoted, flash frozen, then stored at -80ºC. Note that protein used for cryo-EM was processed directly without freezing. LIS1-SNAP required a gel filtration step to improve purity for mass photometry. To this end, the TEV eluate was injected on to a Superdex 200 10/300 equilibrated in TEV buffer (without ATP). Peak fractions were pooled, concentrated, aliquoted, snap frozen, and stored at -80ºC.

### Dynein ATPase assays

ATPase activities were determined using the EnzChek phosphate assay kit (Life Technologies). Assays were performed in motility buffer (30 mM HEPES pH 7.2, 50 mM potassium acetate, 2 mM magnesium acetate, 1 mM EGTA, 1 mM DTT) supplemented with 2 mM MgATP, with or without 2 μM taxol-stabilized microtubules, 5 nM 6His-GST-dynein_331_ (wild-type or mutants). Reactions were initiated with the addition of dynein, and the absorbance at 360 nm was monitored by a spectrophotometer for 10–20 min. Background phosphate release levels (presumably from microtubules) for each reaction were measured for 5 min before addition of dynein to account for any variation as a consequence of differing microtubule concentrations, and were subtracted out from each data point.

### Live cell imaging experiments

Cells were grown to mid-log phase in SD media supplemented with 2% glucose, and mounted on agarose pads. Images were collected at room temperature using a 1.49 NA 100X objective on a Ti-E inverted microscope equipped with a Ti-S-E motorized stage (Nikon), piezo Z-control (Physik Instrumente), a SOLA SM II LE LED light engine (Lumencor), a motorized filter cube turret, and an iXon X3 DU897 cooled EM-CCD camera (Andor). The microscope system was controlled by NIS-Elements software (Nikon). Step sizes of 0.5 µm (for Bik1-3mCherry quantitation) or µm (for dynein quantification) were used to acquire 2-µm-thick Z-stack images. Sputtered/ET filter cube sets (Chroma Technology) were used for imaging mTurquoise2 (49001), GFP (49002), YFP (49003), and mCherry (49008) fluorescence. Images were analyzed using FIJI. Plus end and cortical foci were scored (frequency and intensity) from maximum-intensity projected timelapse movies. Intensity values plotted throughout are background corrected as follows: a 3×3 box drawn around each focus was used to measure signal, while the same size box was drawn around an adjacent region in the cytoplasm to measure background, which was subtracted from the signal.

### Analytical size exclusion chromatography

To assess dynein_MOTOR_-Pac1 binding, equal concentrations of purified motor proteins were first incubated in the indicated nucleotide (3 mM each) for 10 minutes on ice, followed by addition of Pac1. Following a 10 minute incubation on ice, the mixture was injected on to a Superdex 5/150 using an AKTA Pure. Fractions with JFX646-labeled proteins were separated by SDS-PAGE, which were subsequently scanned using a Typhoon gel imaging system (FLA 9500). FIJI was used to determine background-subtracted band intensity.

### Mass photometry

With the exception of LIS1-SNAP, all purified proteins were used directly for mass photometry without additional purification steps. LIS1-SNAP required an additional gel filtration step to remove higher molecular weight species (as determined by mass photometry; see above). All proteins were initially diluted to 500 nM in assay buffer (50 mM Tris, pH 8.0, 150 mM potassium acetate, 2 mM magnesium acetate, 1 mM EGTA, 1 mM DTT) with or without added nucleotide (1 mM of each, as indicated in figures), and then subsequently diluted to 50 nM in same. 3 µl of each was then mixed 1:1 (to 25 nM of each), incubated for 1-2 minutes, and then diluted 1:5 on the stage (2.5 µl of mixed protein + 10 µl same buffer with or without nucleotide) to 5 nM final immediately prior to image acquisition. For apo conditions, residual ATP from the protein preparation was depleted using apyrase by mixing 4.5 µl of 500 nM protein with a 0.5 µl of apyrase (NEB), and incubating for 30 minutes at room temperature. 1 minute movies were acquired using Refeyn MP, and all images were processed and analyzed using Discover MP. Calibration was performed with beta-amylase and thyroglobulin.

### Cryo-EM grid preparation

Purified proteins (as described above) were applied to a TSKgel G4000SWXL column pre-equilibrated with 50 mM Tris pH 7.4, 150 mM potassium acetate, 2 mM magnesium acetate, 1 mM EGTA, 1 mM DTT, and 0.1 mM Mg-ATP. Peak fractions were pooled and 1 mM Mg-ATP with (for MT-U proteins) or without (for MT-B) 1 mM Na_3_VO_4_ was immediately added. Protein quality was examined by negative staining microscopy. Glycerol was added to a final concentration of 10%, aliquots were flash-frozen in liquid nitrogen, and stored at -80 °C. For initial cryo-EM analysis of the human dynein MT-B mutant in ATP buffer (50 mM Tris pH 7.4, 150 mM potassium acetate, 2 mM magnesium acetate, 1 mM EGTA, 1 mM DTT, 1 mM Mg-ATP), we found that a high concentration (> 5 mg/ml) of protein was required for the protein to enter the open holes of a plasma cleaned QUANTIFOIL Au R2/1 300-mesh grids (Fig. S4A). To reduce the sample concentration during cryo-EM grid preparation for human dynein MT-U mutant in ATP-Vi buffer (50 mM Tris pH 7.4, 150 mM potassium acetate, 2 mM magnesium acetate, 1 mM EGTA, 1 mM DTT, 1 mM Mg-ATP and 1 mM Na_3_VO_4_), we coated QUANTIFOIL Cu R2/1 300-mesh grids with graphene-oxide (GO) layers, as previously reported^71^. 4 μl of the MT-U mutant with or without human LIS1 at 0.2 - 0.4 mg/ml were applied to the graphene oxide-coated side of freshly prepared GO-grids (Fig. S4F and K), followed by a 4 s wait time, 3-6 s blot time, 4 blot force, and subsequent freezing in liquid ethane using a Vitrobot Mark IV unit (FEI). The Vitrobot chamber was maintained at close to 95% humidity at 4°C.

The MT-B and MT-U alone data were collected at the Yale ScienceHill-Cryo-EM facility using a Glacios microscope (Thermo Fisher Scientific) operated at 200 keV. The images were collected with a K2 direct electron detector (Gatan) operating in super-resolution mode, at a nominal magnification of 36,000X, corresponding to a pixel size of 1.149 Å. Data collection was automated by SerialEM software^72^ with a defocus range of -1.5 μm to -2.7 μm. In total, 3035 movies for MT-B and 3065 movies for MT-U were collected and each movie was dose-fractionated to 40 frames with a total dose of 40 e-/Å^2^ (Fig. S4, Table S1).

The MT-U + LIS1 data was collected at the Laboratory for BioMolecular Structure (LBMS) using a Titan Krios microscope (Thermo Fisher Scientific) operated at 300 keV and equipped with a K3 detector and a BioQuantum energy filter (Gatan) with a slit width of 15 eV. Data collection was automated by EPU software with a defocus range of -1.5 μm to -2.7 μm, and all movies were recorded in a super-resolution mode at a nominal magnification of 105,000X corresponding to a pixel size of 0.825 Å. Each movie was dose-fractionated to 50 frames with a total dose of 50 e-/Å^2^. A total of 5183 movies were collected (Fig. S4, Table S1).

### Cryo-EM image processing

Cryo-EM data processing workflows are outlined in Figure S4. Recorded movies were pre-processed using cryoSPARC Live^73^ including patch motion correction and patch CTF estimation. Exposures were manually curated and micrographs without graphene oxide were removed.

For the MT-B dataset (Fig. S4A-E), Topaz picker^74^ was used for particle picking. In total, 250,463 particles were extracted with a box size of 360 with a pixel size of 1.149 Å. Multiple rounds of 2D classification were performed to filter the particles. Good particles were used for *ab-initio* reconstruction in cryoSPARC. The reconstructed volume was used for several rounds of heterogenous refinement followed by 2D classification. Finally, 44,752 particles were selected and subjected to non-uniform refinement^75^ followed by two rounds of global and local CTF refinement. A 3.4 Å map was obtained as evaluated using a GSFSC criterion of 0.143.

For the MT-U + LIS1 dataset (Fig. S4F-J), Blob picker in cryoSPARC was used for particle picking. An initial 1,400,918 particles were extracted with a box size 500 and binned to 360 box size, resulting in a pixel size of 1.149 Å. The MT-B map was low-passed to 30 Å and used for heterogenous refinement. After several rounds of heterogenous refinement followed by 2D classification, 182,694 particles were selected and subjected to non-uniform refinement. While the dynein motor region was resolved at better resolution, the LIS1 density appeared to be smeared, suggesting flexibility for LIS1 binding. Before performing 3D classification focusing on LIS1 density, we used a mask around the motor region to perform two rounds of global, local CTF refinement, followed by local refinement to better estimate high-order CTF terms and each particle’s local defocus value. This yielded a 2.8 Å map of the motor region. We then used a mask around the AAA3-AAA5-2 LIS1 density for local 3D classification without alignment in cryoSPARC. After classification, 3 major classed were obtained: 1 LIS1 bound (66,212 particles, 36.2%), “1.5” LIS1 bound (62, 910 particles, 34.4%) and 2 LIS1 bound (53,572 particles, 29.3%; see Fig. S9). The 2 LIS1 bound class was subjected to global refinement (motor with 2 LIS1) and local refinement (AAA3-AAA5 with 2 LIS1), yielding a 3.2 Å and 3.3 Å map, respectively.

For the MT-U alone dataset (Fig. S4K-O), a similar strategy was used. 729,028 particles picked by the Blob picker were extracted with a box size of 360. Multiple rounds of heterogenous and 2D classification were used to clean the particles. Finally, 201,717 particles were subjected to non-uniform refinement. Two rounds of global and local CTF refinement followed by local refinement allowed us to obtain a 2.9 Å map.

Local resolution estimation of all maps was performed in cryoSPARC. Directional anisotropy analysis of all maps was performed using 3DFSC^76^ implemented in cryoSPARC.

### Model building and refinement

A previously reported human dynein motor structure^7^ (PDB: 5NUG) was used as an initial model. Individual domains (linker, AAA large, AAA small) were docked into the cryo-EM map as rigid bodies using UCSF ChimeraX^77^. After docking, Namdinator^78^, a molecular dynamics flexible fitting tool, was used to further fit the model into the cryo-EM map. The model was then manually inspected and adjusted in COOT v0.9.5^79,80^. The high-resolution cryo-EM map together with our biochemical assay allowed us to confidently assign nucleotides to each pocket. For example, by adjusting the contour level, we could see the separation of vanadate, Mg^2+^, and ADP in the MTU AAA1 pocket. Our vanadate-mediated photocleavage assay also indicated that vanadate binds to the MT-U AAA1 pocket. These two pieces of evidence allowed us to build ADP-Vi into the MTU AAA1 pocket.

To build the model for human LIS1, the predicted structure from AlphaFold^81^ database was used as the initial model. The positions of two LIS1 were determined using a previously reported yeast dynein-Pac1 structure^55^ (PDB: 7MGM). All models were iteratively refined using Phenix real-space refinement 1.19.2_4158^82^ and manual rebuilding in COOT. The quality of the refined model was analyzed by MolProbity integrated into Phenix^83^ with statistics reported in Table S1. Figures were prepared using UCSF ChimeraX^77^.

### Molecular dynamics simulation and interface energy calculations

The cryo-EM structure of the MT-U-2 LIS1 complex was prepared before modeling and simulations in Charmm-GUI^84^. The large-scale Atomic/Molecular Massively Parallel Simulator was applied for the simulations^85^. Periodic boundary conditions were used to produce a series of proteins. Amber10:EHT force field (https://infoscience.epfl.ch/record/121435/files/Amber10i.pdf) was used to simulate proteins. Water molecules were simulated using the rigid SPC/E force field^86^ whereas the SHAKE algorithm^87^ was used to keep the water molecules rigid throughout the entire simulation. Lennard-Jones 12-6 term^88^ is used to describe the short-range interactions and the cutoff distance was 12 Å. The particle-particle/particle-mesh method with a precision value of 10^−6^ was adopted to estimate long-range electrostatic interactions^89^. First, we ran the energy minimization for the whole system. Next, the simulations were carried out at 25ºC using a canonical NVT ensemble, where the temperature was controlled by the Nosé-Hoover thermostat^90^. Then NPT ensemble was performed in production phase where the target pressure and temperature were 1 atm and 25ºC respectively. Default tether restraints from LAMMPS were applied to the system.

Protein models after in silico mutations underwent the same preparation procedure. Interface energy was calculated in the production phase. The interface energy calculation between contacting residue pairs was processed. The proximity threshold was set to 12 Å. Atoms separated by more than this distance were not considered to be interacting. Six types of contacts were identified: hydrogen bonds, metal, ionic, arene, covalent and Van der Waals distance interactions.

## SUPPLEMENTAL INFORMATION

**Figure S1.**
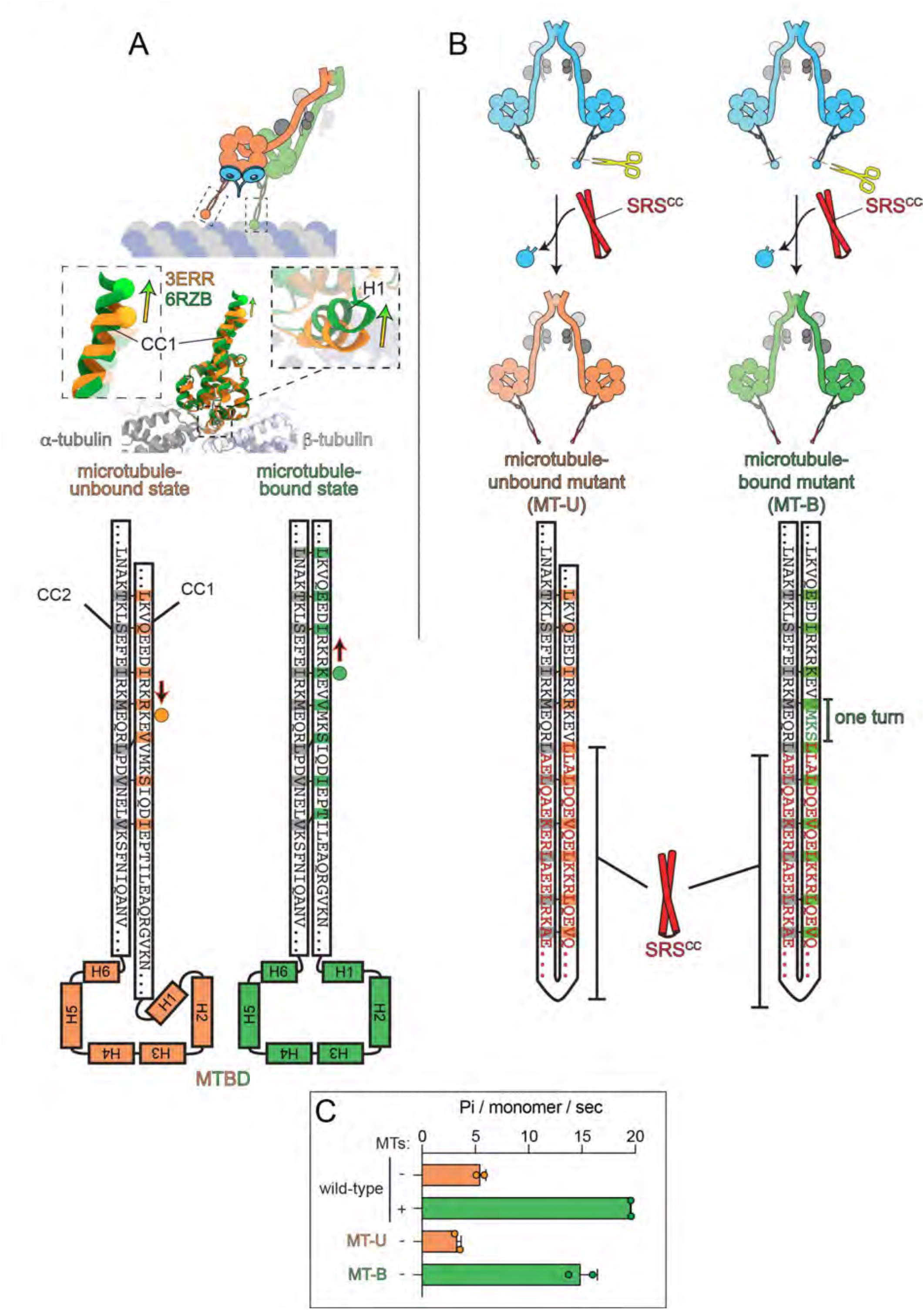
Design strategy and validation of microtubule-bound and -unbound dynein mutants. (A) Cartoon and structural depiction of conformational change that takes place at the coiled-coil (CC) stalk and microtubule-binding domain (MTBD) upon microtubule binding. Comparison of a high resolution microtubule-bound dynein MTBD (6RZB)^32^ and a crystal structure of a microtubule unbound MTBD (3ERR)^34^ reveals an upward shifting of helix 1 (H1) as a result of microtubule binding. This results in a consequent upward shift of CC1 with respect to CC2. (B) Design strategy to generate constitutive microtubule-unbound and -bound dynein mutants. The stable coiled-coil from seryl tRNA synthetase (SRS^CC^) was used to replace the entire dynein MTBD plus short regions of CC1 and CC2. The MT-B mutant has 4 additional residues in CC1 with respect to the MT-U mutant (corresponding to one helix turn), resulting in a presumed upward shift of CC1 compared to CC2. (C) Plot depicting ATPase levels for artificially dimerized dynein motor domain fragments (GST-dynein_MOTOR_). Note the MT-U mutant closely reflects the wild-type dynein motor in the absence of microtubules, while the MT-B mutant matches that of wild-type plus a saturating concentration of microtubules (2 µM)^70^.

**Figure S2.**
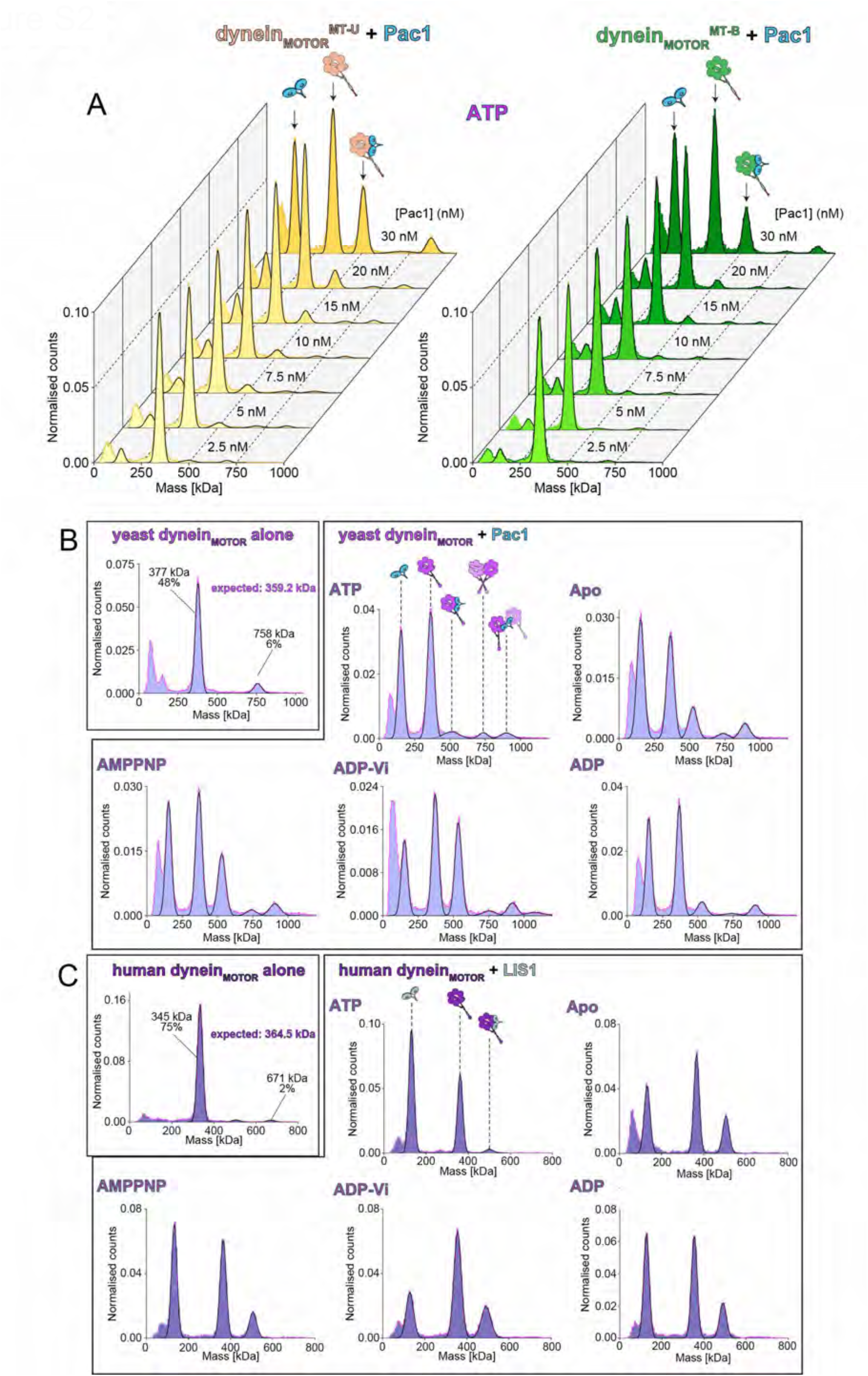
Additional mass photometric analysis of Pac1- and LIS1-dynein_MOTOR_ binding. (A) Histograms of mass photometry analysis depicting relative dynein_MOTOR_-Pac1 complex formation in the presence of a fixed concentration of each motor (25 nM) and increasing concentrations of Pac1 (as indicated). Note the higher fraction of dynein_MOTOR_^MT-U^-Pac1 complex formation with respect to dynein_MOTOR_^MT-B^-Pac1 at every concentration. (B and C) Representative mass histograms for the wild-type yeast (B) and human (C) dynein_MOTOR_ proteins with and without Pac1 or LIS1 with different nucleotides, as indicated (see Figure 3 and Methods). Cartoon depictions above each peak in the ATP panel indicates the likely dynein_MOTOR_-Pac1 or LIS1 complex contained therein.

**Figure S3.**
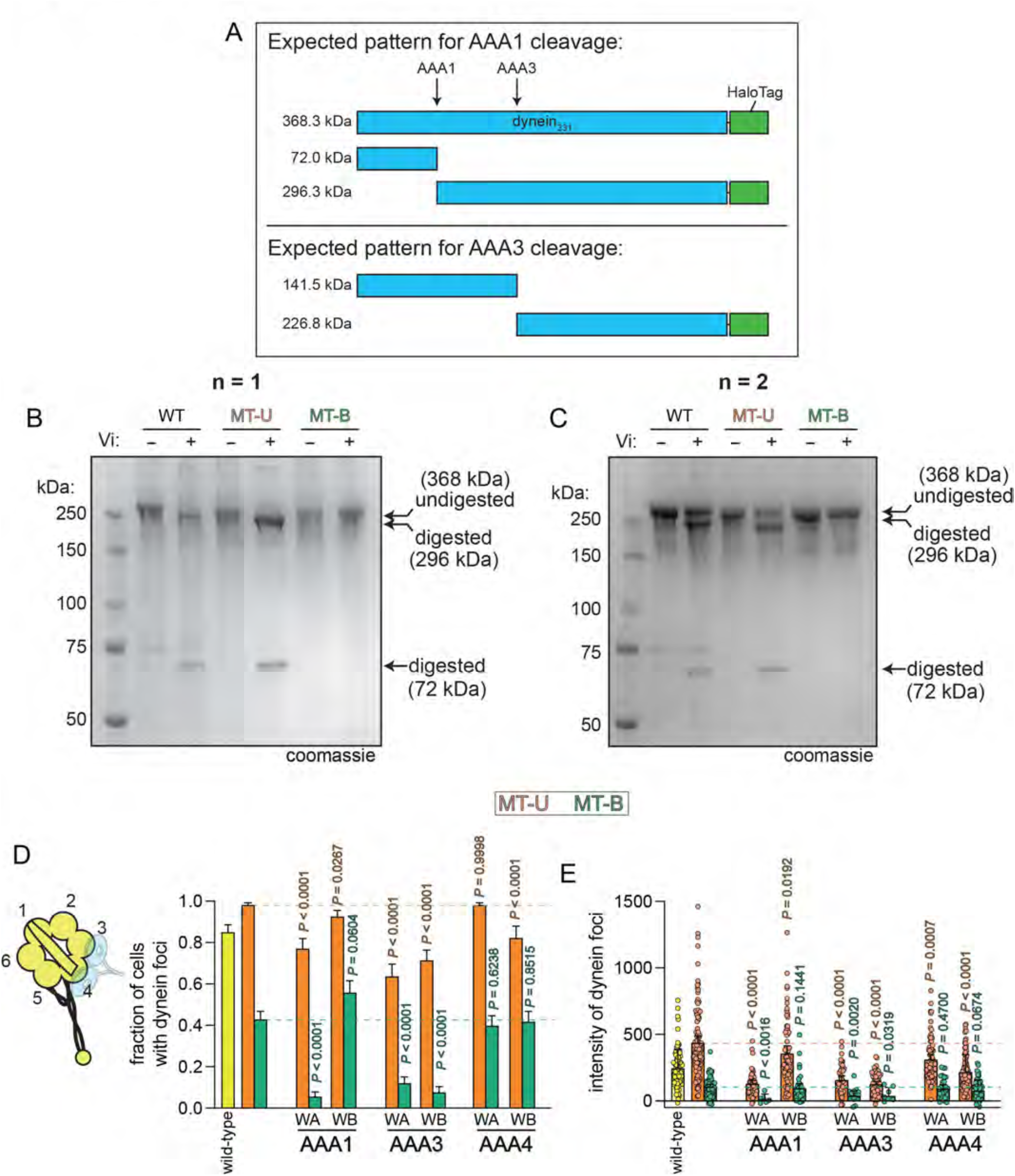
Vanadate-mediated photocleavage assay, and the role of ATP binding and hydrolysis in dynein-Pac1 binding. (A) Schematic depicting expected vanadate-mediated photocleavage if vanadate were bound to AAA1 (top) or AAA3 (bottom). (B and C) Two independent replicates of photocleavage assay. Purified indicated dynein_MOTOR_ fragments were incubated with 3 mM ATP with or without 3 mM vanadate, exposed to ultraviolet light (254 nm) for 1 hour, and then analyzed by coomassie-stained SDS-PAGE. (D) Plots depicting frequency and intensity of dynein foci in cells expressing indicated *DYN1* allele as the only source of dynein heavy chain (“WA”, Walker A mutant; “WB”, Walker B mutant). Mutations in Walker motifs are as follows: AAA1 WA, K1802A; AAA1 WB, E1849Q; AAA3 WA, K2424A; AAA3 WB, E2488Q; AAA4 WA, K2766A; AAA4 WB, E2819Q. Foci were scored from timelapse movies. *P* values indicate statistical significance of data sets (calculated using a Mann-Whitney test) with respect to wild-type dynein_MOTOR_^MT-U^or dynein_MOTOR_^MT-B^.

**Figure S4.**
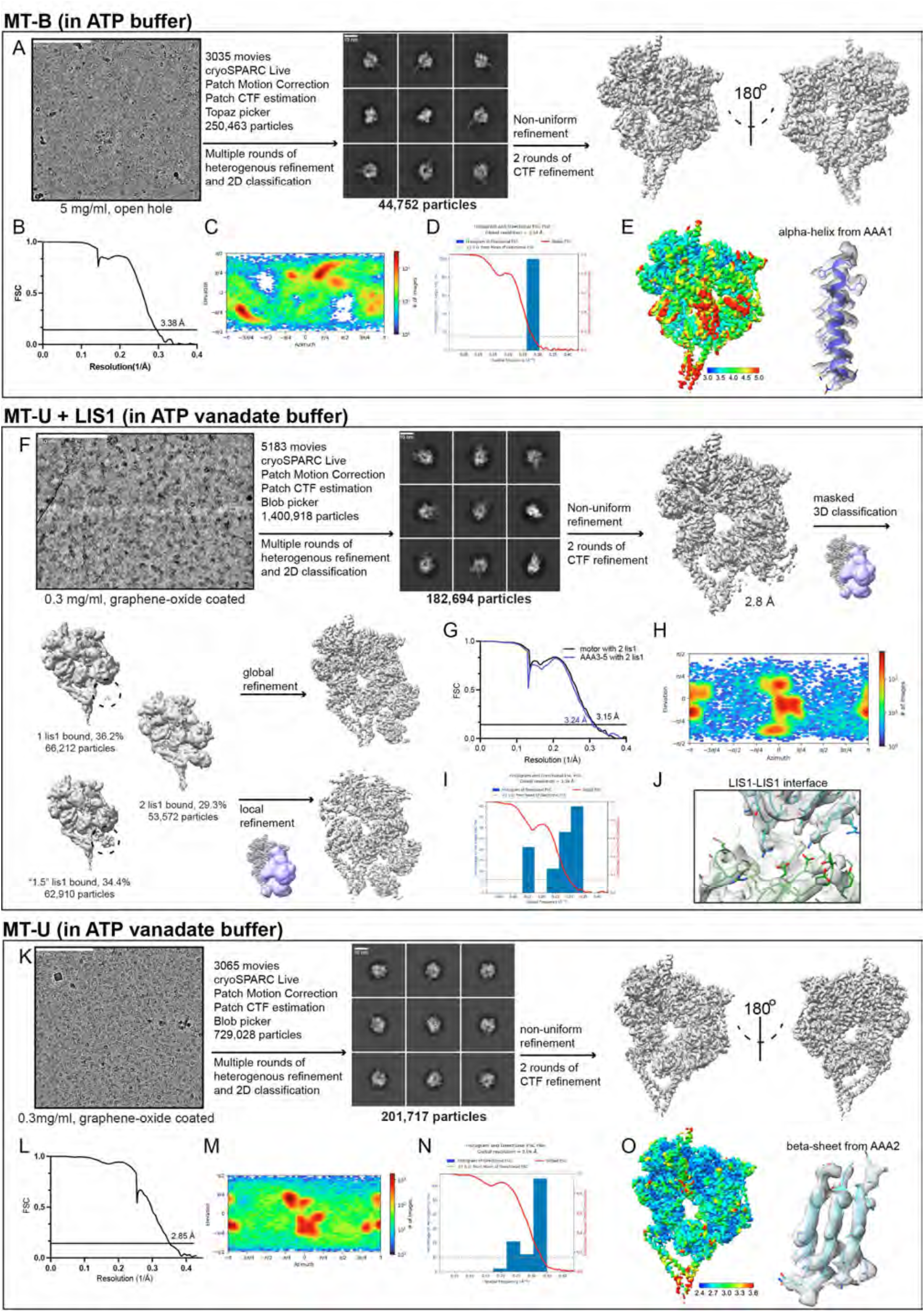
Cryo-EM data processing flowchart. (A) Cryo-EM workflow of MT-B in the presence of ATP (details in methods). (B) FSC curves with the gold standard threshold of 0.143 for MT-B. (C - D) Particle distribution plot and 3D FSC analysis of MT-B. (E) Local resolution analysis of MT-B and representative cryo-EM densities. (F) Cryo-EM workflow of MT-U with LIS1. (G) FSC curves with the gold standard threshold of 0.143. (H - I) Particle distribution plot and 3D FSC analysis of MT-U + LIS1 in the presence of ATP-Vi. (J) Local resolution analysis and representative cryo-EM densities of the LIS1-LIS1 interface. (K) Cryo-EM workflow of MT-U alone in the presence of ATP-Vi. (L) FSC curves with the gold standard threshold of 0.143. (M - N) Particle distribution plot and 3D FSC analysis of MT-U. (O) Local resolution analysis and representative cryo-EM densities.

**Figure S5.**
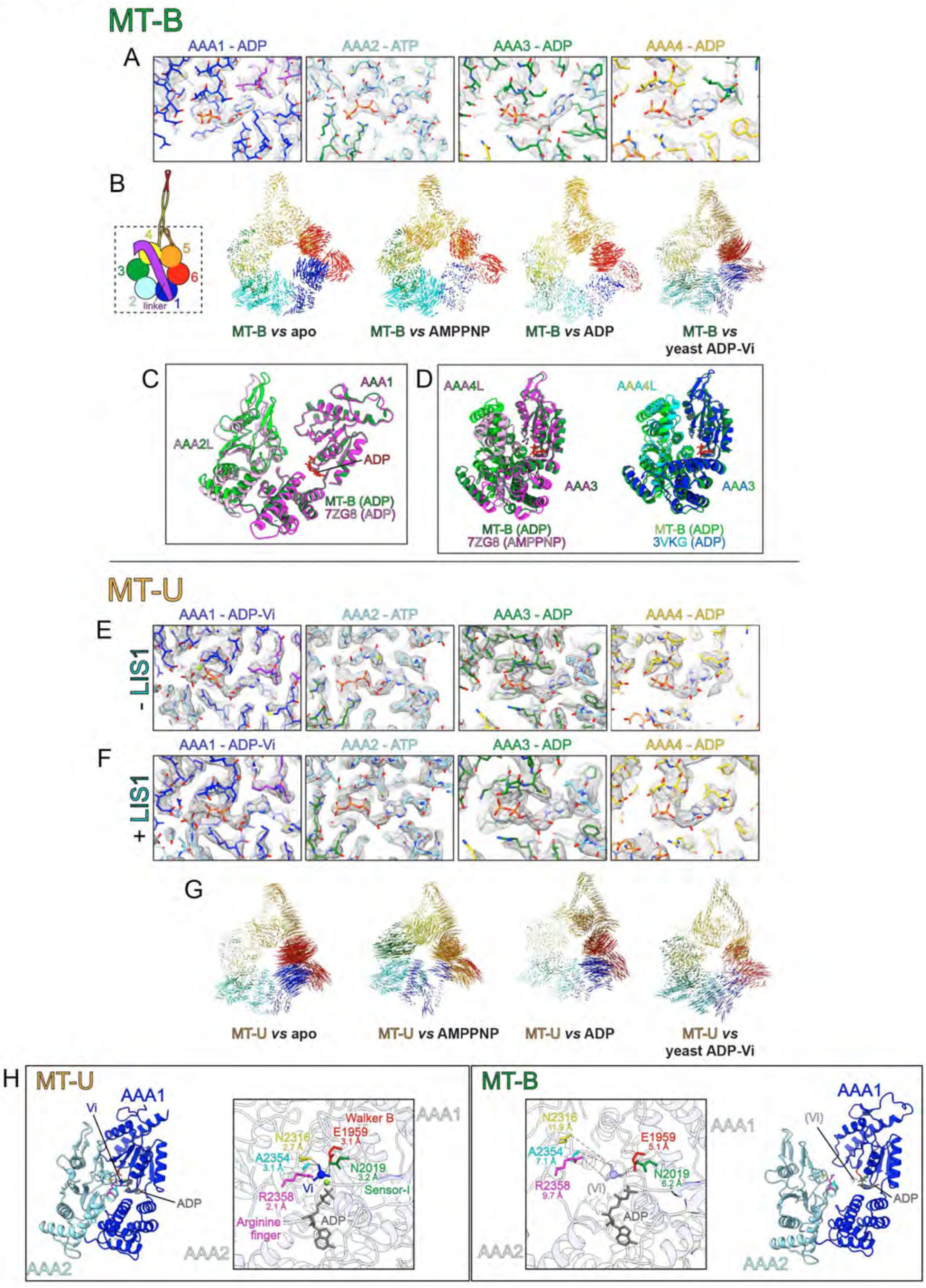
Additional analysis of human MT-U and MT-B cryo-EM structures. (A) Stick representations of the dynein_MOTOR_^MT-B^AAA sites showing the nucleotide electron density (the center of each image) and surrounding residues (residues are color-coded as shown in panel B schematic). (B) Vector maps depicting pairwise alpha carbon interatomic distances between the dynein_MOTOR_^MT-B^ with the following: apo yeast dynein (4AKG), AMPPNP-bound yeast dynein (4W8F), ADP-bound *Dictyostelium discoideum* dynein (3VKG), ADP-Vi and Pac1-bound yeast dynein (7MGM)^41,52,63,91^. Structures were globally aligned after removal of the linkers. (C) AAA1-AAA2L domains from dynein_MOTOR_^MT-U^ (shades of green) and the native microtubule-bound dynein-1 (magenta and pink) overlaid to depict the high degree of structural similarity. The two were locally aligned using AAA1. (D) AAA3-AAA4L domains from dynein_MOTOR_^MT-U^ (shades of green) overlaid with either the native microtubule-bound dynein-1 (left, magenta and pink) or the ADP-bound *Dictyostelium discoideum* dynein (right, blue and green). (E and F) Stick representations of the LIS1-unbound (E) or bound (F) dynein_MOTOR_^MT-U^AAA sites showing the nucleotide electron density and surrounding residues (residues are color-coded as shown in panel B schematic). (G) Vector maps depicting pairwise alpha carbon interatomic distances between the LIS1-bound dynein_MOTOR_^MT-U^ with those described for panel B. Structures were globally aligned after removal of the linkers. (H) AAA1-AAA2L domains from dynein_MOTOR_^MT-U^ (left) and dynein_MOTOR_^MT-B^ (right) with residues of dynein_MOTOR_^MT-U^ contacting the Vi highlighted (E1959, Walker B; N2019, Sensor-I; R2358, arginine finger; N2316; A2354;). Distances between these residues and the Vi (or, for the dynein_MOTOR_^MT-B^ structure, between them and where the Vi would be) are indicated.

**Figure S6.**
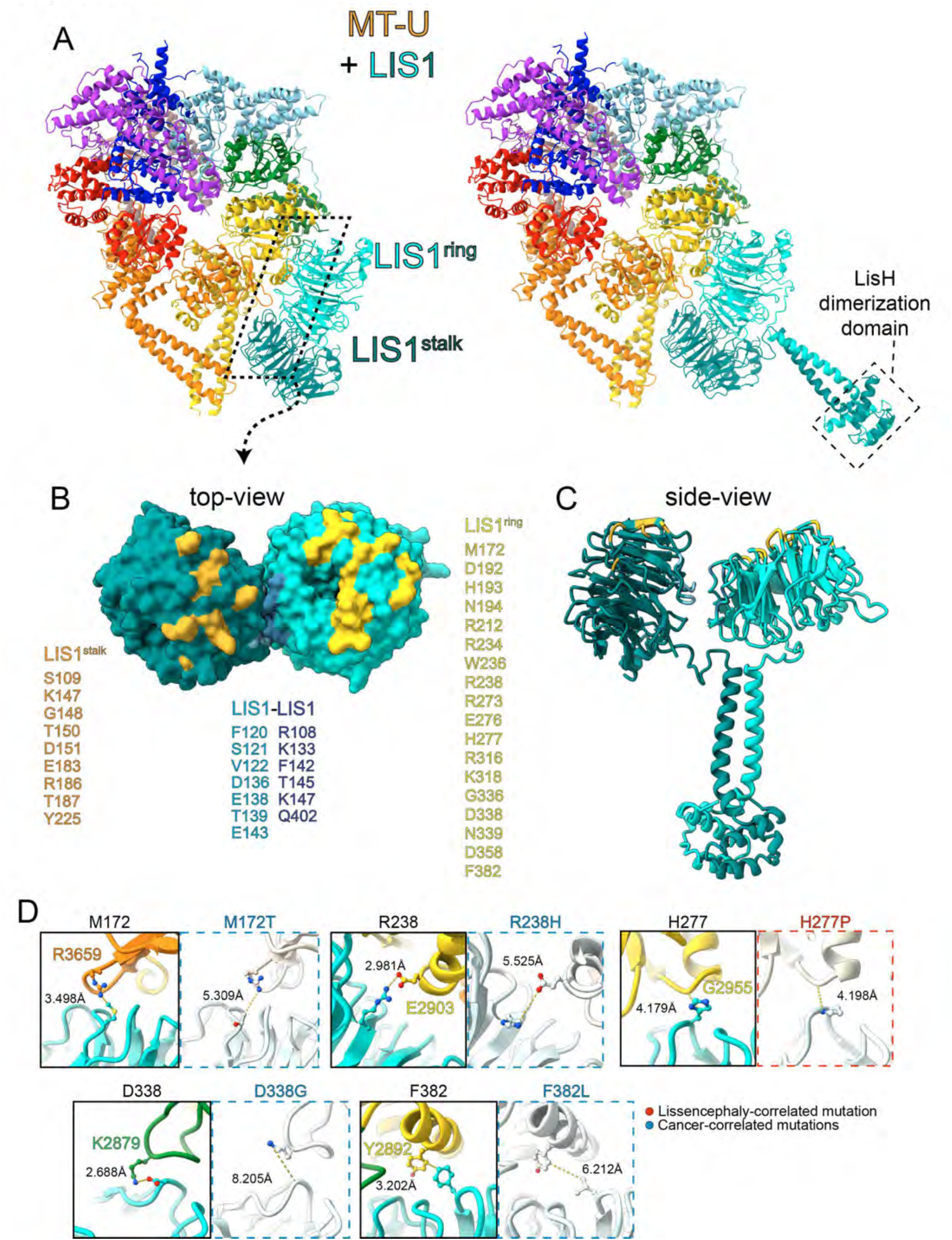
Additional analysis of LIS1-bound dynein_MOTOR_^MT-U^ structure. (A) 2 LIS1-bound dynein_MOTOR_^MT-U^ structure with domains colored as shown in Figure 5 (left) and the same shown with the a full-length LIS1 homodimer, with the N-terminal dimerization domain modeled. The LIS1 N-terminal dimer model was generated using a combination of AlphaFold prediction^81^ and a previous crystal structure, 1UUJ^92^. The structure was manually adjusted in COOT. (B) View of LIS1 homodimer surface that contacts site^stalk^ (teal) and site^ring^ (cyan). Residues listed and indicated in different colors on the structure are those that make direct contact with dynein or LIS1. (C) Side-view of full-length LIS1 homodimer model with residues colored as in panel B. (D) Results of molecular dynamics simulation from Figure 5G depicting change in interatomic distances in residues mediating contacts between LIS1 and dynein as a consequence of indicated mutations.

**Figure S7.**
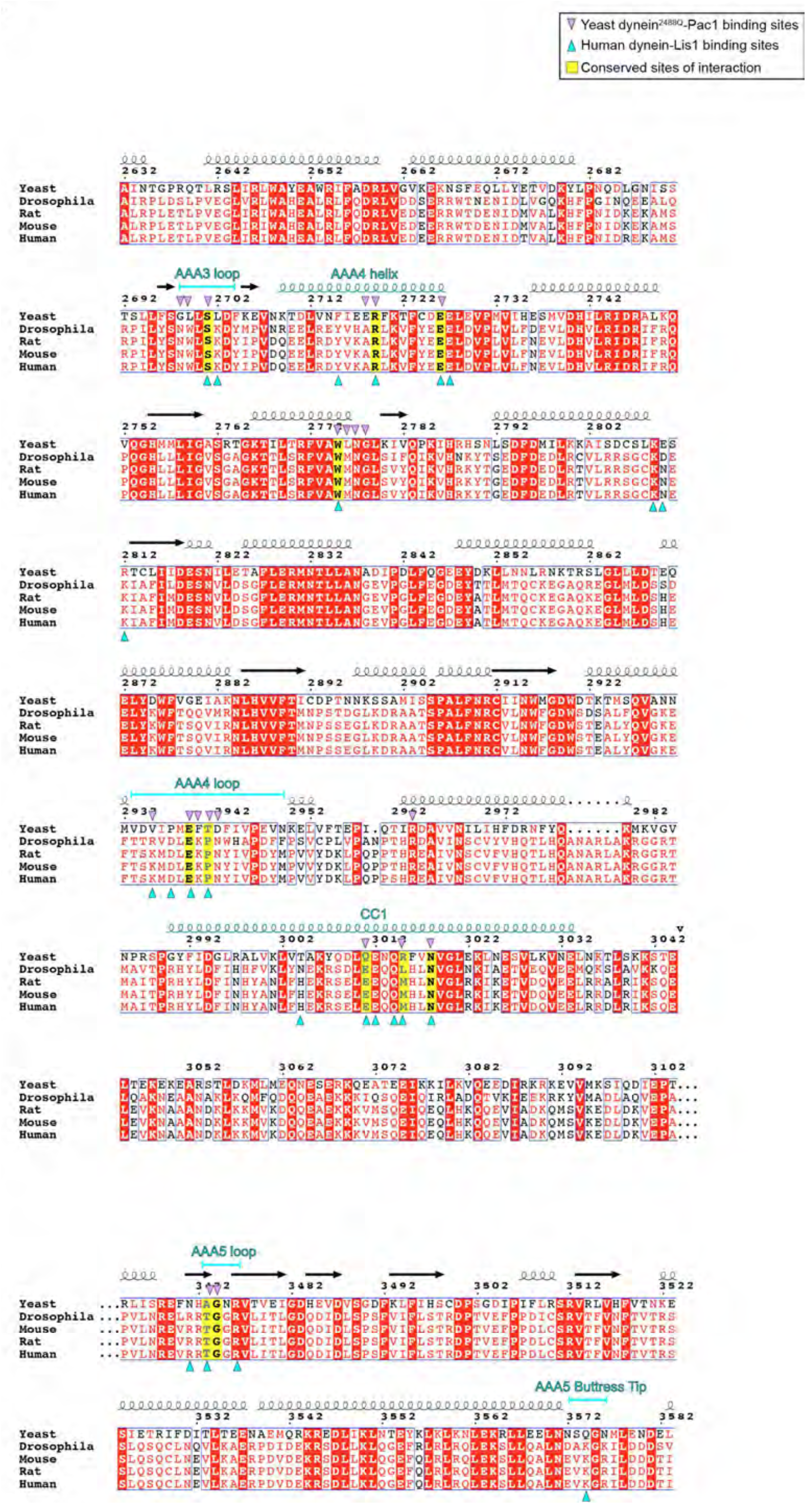
Sequence alignment of Pac1 and LIS1-binding regions within dynein. Numbering corresponds to yeast dynein (Dyn1) sequence. Secondary structure indicated with cartoon helices (for alpha-helices) and arrows (for beta-strands).

**Figure S8.**
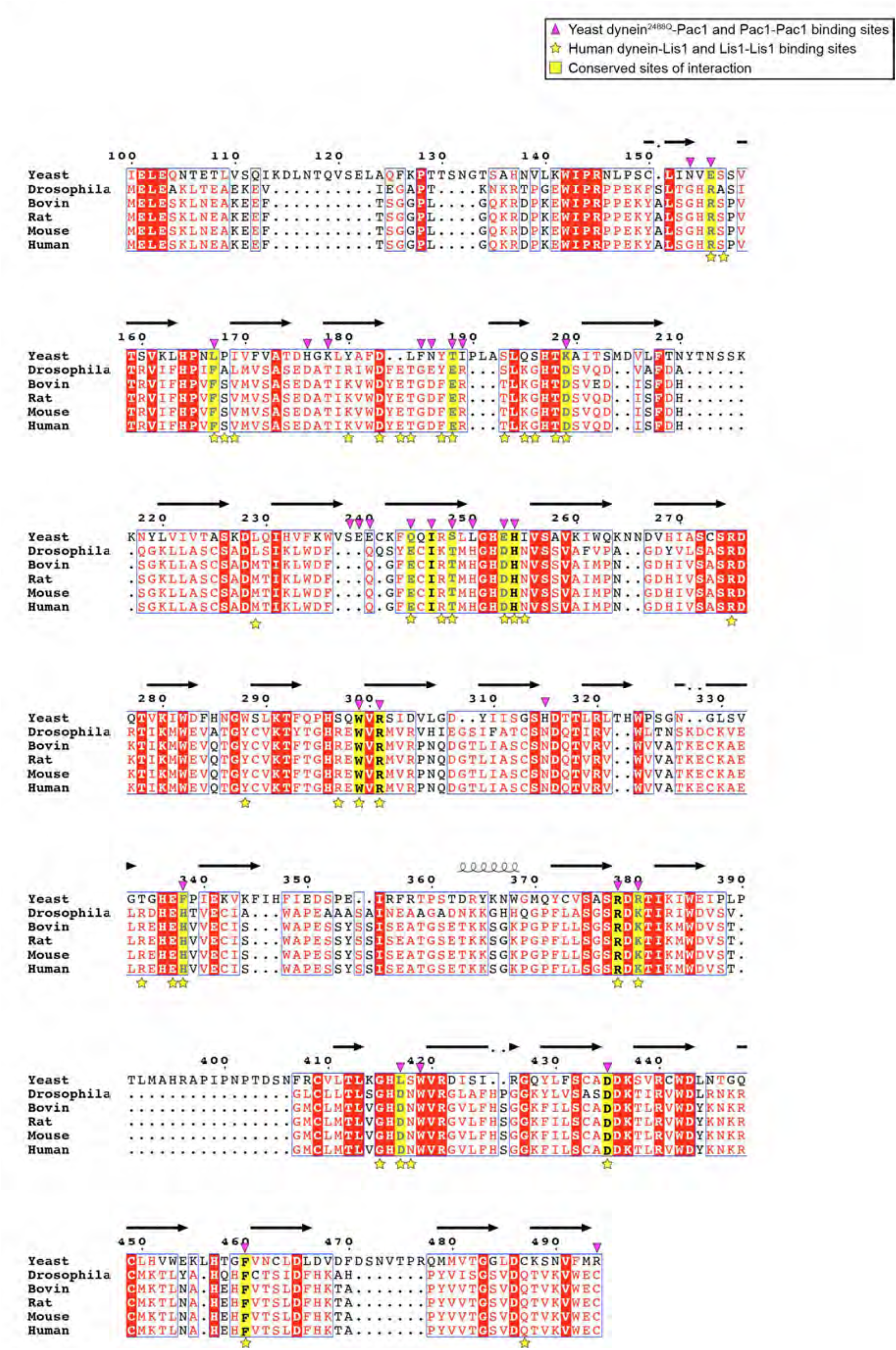
Sequence alignment of dynein-binding regions within LIS1 and homologs. Numbering corresponds to yeast Pac1 sequence. Secondary structure indicated with cartoon helices (for alpha-helices) and arrows (for beta-strands).

**Figure S9.**
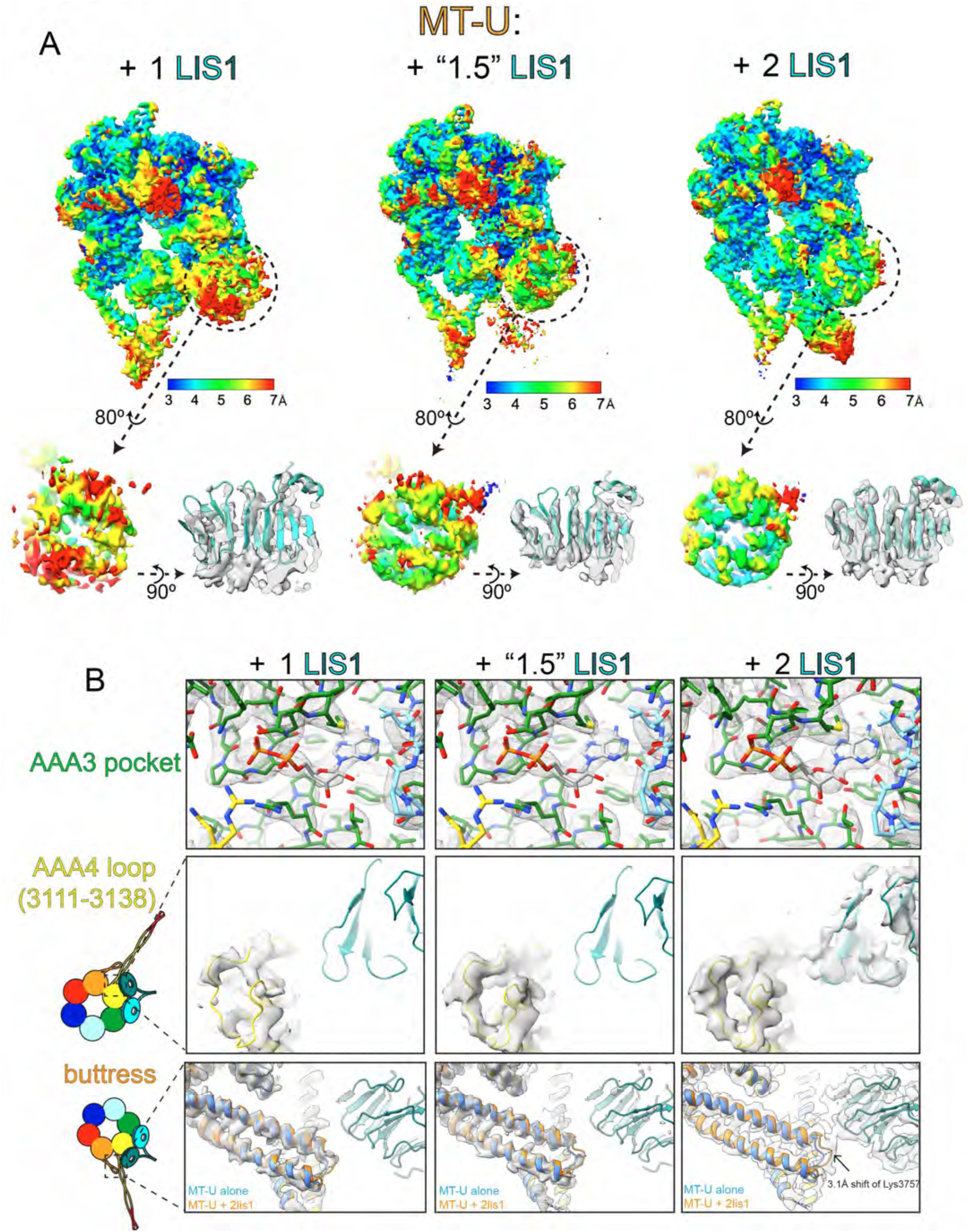
3D classification analysis of LIS1-bound dynein_MOTOR_^MT-U^ structures. (A) The three classes of LIS1-bound dynein (shown with a rotated close-up view of LIS1^ring^) are shown with local resolution indicated by color. Note the significant increase in resolution and map quality for LIS1^ring^ for the “1.5” and LIS1-bound dyneins. (B) Close-up views of the indicated regions of the indicated LIS1-bound dynein_MOTOR_^MT-U^ structure. Note all three structures have clear density for ADP at AAA3 (top), and the improvement in local resolution for the AAA4 loop (middle) and the buttress for the “1.5” and 2 LIS1-bound structures.

## SUPPLEMENTAL VIDEO LEGENDS

Video S1. **Examples of Bik1-labeled astral microtubules with plus ends statically associated with cortex**. Timelapse fluorescence images of cells expressing Bik1-3mCherry, GFP-Tub1, and Dyn1^MT-U^-3YFP (arrow in first frame on merge, astral microtubule plus ends with Bik1 foci statically associated with the cortex). Cartoons represent cells in first frame of movie (small green circle, spindle pole body; green lines, astral microtubules; magenta circles, plus end Bik1 foci statically associated with cortex).

Video S2. **Close-up views of dynein**^**MT-U**^**-LIS1 contacts**. Intermolecular contacts between dynein and LIS1, and between LIS1 and LIS1, are indicated with dashed lines. Nucleotide densities in AAA1 and AAA3 are indicated with mesh outlines.

Video S3. **Conformational differences within the AAA+ ring between MT-U and MT-B dynein**. Structures for dynein_MOTOR_^MT-U^ (+ LIS1) and dynein_MOTOR_^MT-B^ were globally aligned (after removal of the linker and C-terminal domains). ChimeraX was used to morph between these two structures. Inset highlights the changes in AAA4-6 that appear to be initiated by the bending of the buttress. Note the rigid body reorientation of AAA5S-AAA6L, which transmits structural rearrangements to AAA6S and AAA1.

Video S4. **Conformational differences at the LIS1-binding sites between MT-U and MT-B dynein**. Structures for dynein_MOTOR_^MT-U^ (+ LIS1) and dynein_MOTOR_^MT-U^were locally aligned using AAA4-AAA5. ChimeraX was used to morph between these two structures. Shown are closeup views of site^stalk^ (left) and site^ring^ (right). Residues with sidechains shown on LIS1 and dynein are those that mediate intermolecular contacts.

Video S5. **Model for Pac1/LIS1 function in budding yeast**. See text for detailed description of model.

**Table S1.**
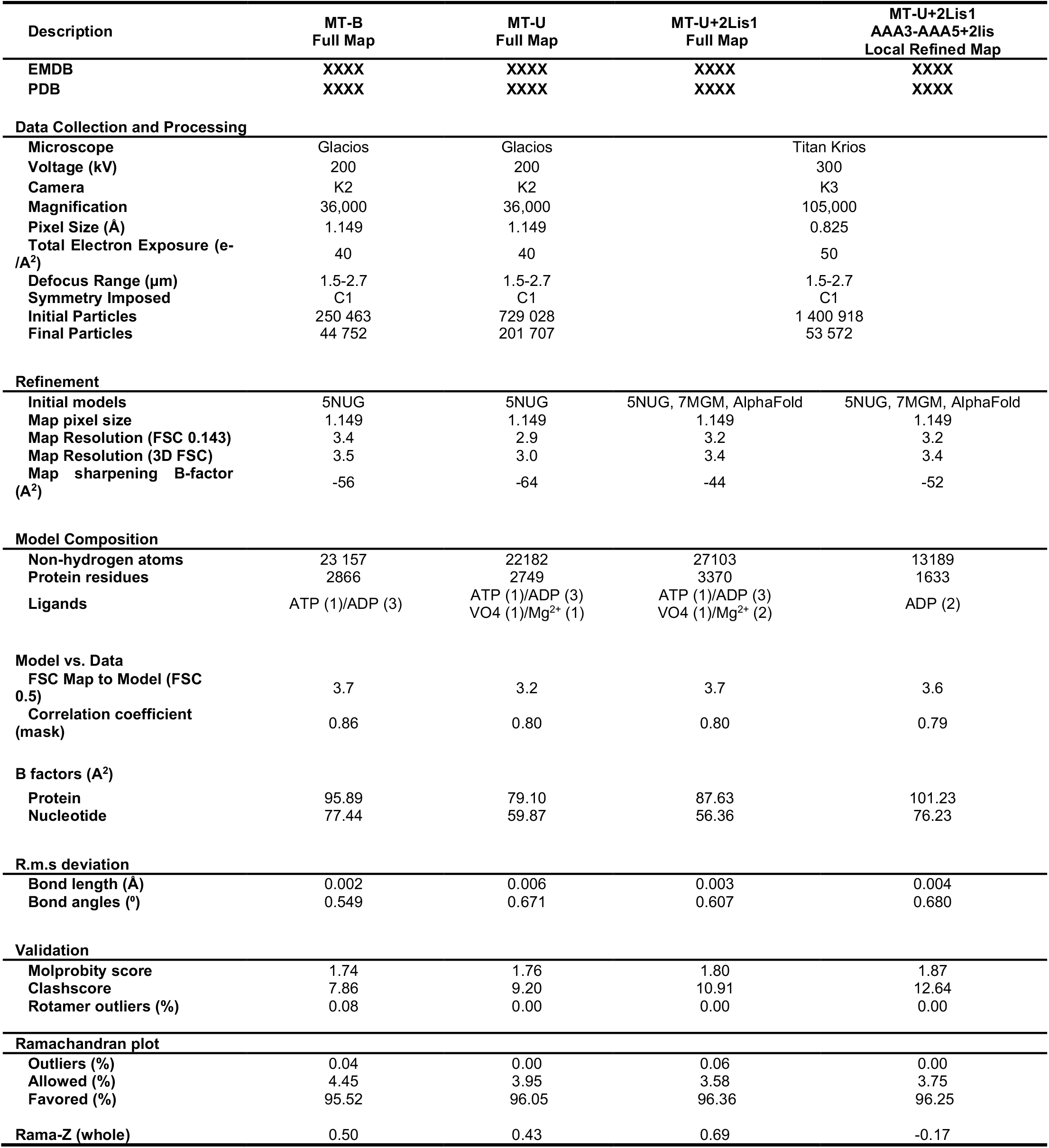
Cryo-EM Data Collection, Refinement, and Validation Statistics.

